# PlasGO: enhancing GO-based function prediction for plasmid-encoded proteins based on genetic structure

**DOI:** 10.1101/2024.07.03.602011

**Authors:** Yongxin Ji, Jiayu Shang, Jiaojiao Guan, Wei Zou, Herui Liao, Xubo Tang, Yanni Sun

## Abstract

Plasmid, as a mobile genetic element, plays a pivotal role in facilitating the transfer of traits, such as antimicrobial resistance, among the bacterial community. Annotating plasmid-encoded proteins with the widely used Gene Ontology (GO) vocabulary is a fundamental step in various tasks, including plasmid mobility classification. However, GO prediction for plasmid-encoded proteins faces two major challenges: the high diversity of functions and the limited availability of high-quality GO annotations. Thus, we introduce PlasGO, a tool that leverages a hierarchical architecture to predict GO terms for plasmid proteins. PlasGO utilizes a powerful protein language model to learn the local context within protein sentences and a BERT model to capture the global context within plasmid sentences. Additionally, PlasGO allows users to control the precision by incorporating a self-attention confidence weighting mechanism. We rigorously evaluated PlasGO and benchmarked it against six state-of-the-art tools in a series of experiments. The experimental results collectively demonstrate that PlasGO has achieved commendable performance. PlasGO significantly expanded the annotations of the plasmid-encoded protein database by assigning high-confidence GO terms to over 95% of previously unannotated proteins, showcasing impressive precision of 0.8229, 0.7941, and 0.8870 for the three GO categories, respectively, as measured on the novel protein test set.

## Introduction

Plasmids are typically circular, extrachromosomal DNA molecules primarily found in bacteria. As a type of mobile genetic element (MGE), about half of them can mediate horizontal gene transfer (HGT) by transferring between different bacteria through a process known as conjugation [1, 2]. Consequently, the advantageous traits carried by the conjugative plasmids, such as antimicrobial resistance (AMR), will be disseminated among the bacterial community, thereby promoting the evolutionary adaptation of the host bacteria [3]. Plasmid-specific proteins can be classified into two main categories: core (backbone) proteins and accessory (payload) proteins [4]. The core proteins play essential roles in plasmids’ housekeeping functions, encompassing replication, stability, and conjugation. They are instrumental in plasmid typing, such as replicon (Rep) typing, mobilization (MOB) typing, and mate-pair formation (MPF) typing [5]. Moreover, the core proteins exhibit a strong correlation with the plasmid host range, namely the hosts to which the plasmid can be transferred or in which it can be maintained. On the other hand, the accessory proteins encode the host-beneficial traits, such as AMR and virulence. In summary, functional annotation of plasmid-encoded proteins not only helps identify the plasmid-carried traits but also provides valuable insights into predicting the transmission trajectory of these traits.

Gene Ontology (GO) is one of the most widely adopted vocabularies for describing protein functions, encompassing a collection of 42,255 GO terms by April 2024 [6]. The GO terms are structured in a directed acyclic graph (DAG) format, with three root terms representing the three GO categories: Molecular Function (MF), Biological Process (BP), and Cellular Component (CC). Therefore, a protein can be annotated with GO terms from all three aspects to achieve a comprehensive description of its functionality. According to the true path rule, if a protein is annotated with a specific GO term, it is also considered to be annotated with all the ancestor terms (semantically more general) of that particular GO term. Therefore, a protein can be annotated with more than one GO term, leading to the formulation of GO term prediction as a multi-label classification problem.

The rapid development of deep learning (DL) offers a promising approach, leveraging its strong generalization capability to predict the functions of novel proteins. Several DL methods have been proposed for GO term prediction utilizing only protein sequences as input. Among them, DeepSeq [7] and DeepGOPlus [8] are designed based on convolutional neural networks (CNNs). Specifically, DeepSeq first utilizes word embedding to represent each amino acid (AA) as a 23-dimensional vector and then feeds these embeddings to two 1D convolutional (Conv1d) layers to extract meaningful features for function prediction. DeepGOPlus enhances the performance by incorporating 16 Conv1d layers with varying filter sizes, ranging from 8 to 128, in parallel. This enables the model to learn sequence motifs of different lengths, which play a crucial role in predicting protein function. The other two methods, TALE [9], and PFresGO [10], are built on the Transformer architecture and leverage the hierarchical information in the GO DAG. While using a Transformer layer to capture the dependencies among AAs within a protein, TALE also learns embeddings for the GO term labels to encode the ancestor relationship between GO terms. By multiplying the features learned by the Transformer and the label embeddings, a similarity matrix is generated for the final prediction, representing the similarity score between each AA and each label. PFresGO replaces the self-attention model in Transformer with a cross-attention mechanism, where GO terms act as the query to identify the functionally relevant AAs. Although there are a number of GO term prediction tools, they are not optimized for plasmid-encoded proteins.

Challenges for plasmid protein GO prediction

The prediction of GO terms for plasmid-encoded proteins presents two major challenges that have not been well-addressed by generic GO prediction tools. First, compared to other types of biological entities, plasmids tend to encode a smaller number of proteins, but these proteins showcase a comparable level of functional diversity to more complicated peers. For instance, the manually reviewed Swiss-Prot database includes 323,202 proteins encoded in bacterial chromosomes, associated with 9,631 GO terms. In contrast, plasmid-encoded proteins, despite only accounting for nearly one percent of the protein count (3,202), are still associated with a considerable number of GO terms (3,318). This larger ratio of GO term to protein count can be attributed to two main causes: the frequent genetic exchange events occurring between chromosomes and plasmids [11], as well as the gene flow between plasmids and phages mediated by phage-plasmid elements [12]. As a result, plasmids encode many proteins derived from both chromosomes and phages. Overall, the combination of a large number of GO terms (labels) and a relatively small number of proteins (training samples) increases the difficulty of the multi-label classification. The second challenge relates to the limited availability of high-quality GO annotations for plasmid-encoded proteins. For instance, considering the 678,197 non-redundant proteins encoded in 47,871 complete plasmids obtained from the RefSeq database, only 29.34% of these proteins possess annotated GO terms from at least one of the three GO categories. Alignment-based methods like the InterPro2GO pipeline [13] failed to extend the protein annotation rate due to their inability to identify signatures (protein families or domains) for the remaining uncharacterized proteins.

To address these challenges, we design a method that capitalizes on the genetic structures of plasmids for better GO prediction. Like human languages that possess a linguistic structure, genes residing on plasmids also exhibit a distinct biological structure, characterized by a modularization pattern [4]. This pattern often results in the division of a plasmid into functionally related segments, including areas dedicated to replication, conjugation, payload, and other plasmid-specific functions. The reason for this phenomenon is the dynamic evolution of plasmids over time, facilitated by the acquisition or loss of segments through recombination or the movement of MGEs [14]. Furthermore, functionally related segments within plasmids resemble phrases in human languages, attributable to both the similarity of protein functions and specific interactions between certain proteins within the same segment (e.g., relaxases interacting with type IV coupling proteins during conjugation). As depicted in Figure 1, the large conjugative multidrug resistance plasmid pOLA52 [15] explicitly displays a modularization pattern, wherein the coding sequences (CDSs) with similar functions are more likely to be closely positioned. Interestingly, as shown on the right side of Figure 1, several payload genes are interspersed with two transposases (a type of MGE gene), suggesting that this specific segment may have originated from HGT facilitated by transposons. This type of modular structure can be learned by natural language models, which will be the major component in our methodology.

**Figure 1.**
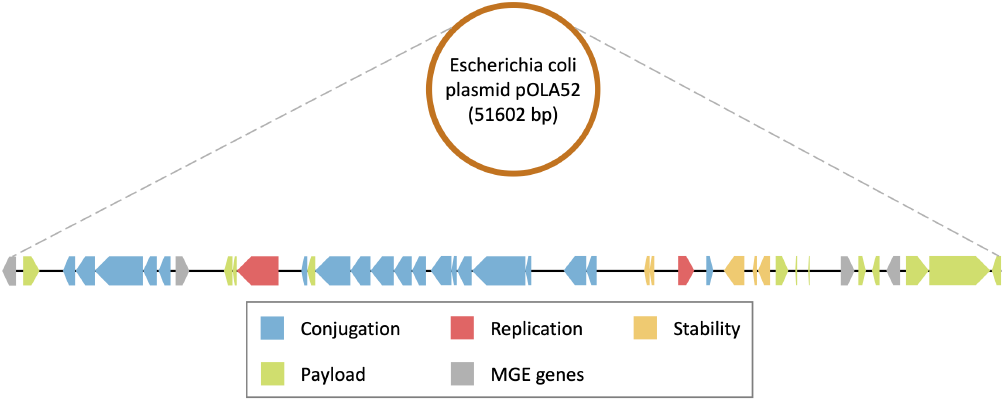
The flattened diagram of the circular plasmid pOLA52, which illustrates the CDS region of each gene, with the color representing the function of the encoded protein. The encoded proteins were manually classified into five functional classes based on the gene product annotation in the NCBI database. In the diagram, a reversed pentagon block indicates that the corresponding CDS is located on the complementary strand. Additionally, CDSs without gene product annotation were excluded from this diagram.

In addition, we will leverage protein language models (**PLM**s), which have demonstrated remarkable performance in various protein-related tasks by understanding the underlying semantic meaning of the language of life [16]. After self-supervised pre-training on a large corpus of protein sequences without manual labeling, foundation PLMs can generate protein embeddings that encapsulate learned knowledge. An important example of such knowledge is the correlation between one-dimensional (1D) protein sequences and their corresponding three-dimensional (3D) structures. The acquired prior biological knowledge can be leveraged to enhance protein function prediction through transfer learning. As an example, the language model CaLM [17] represents 64 codons as separate tokens and is trained using an unsupervised masked language modeling (MLM) objective. Experimental results demonstrate that the protein representations derived from CaLM outperform other PLMs in the classification of GO terms.

In this study, we have incorporated the above key observations and developed a dedicated tool called PlasGO for predicting GO terms of plasmid-encoded proteins. Rather than starting from scratch, we employ the state-of-the-art (SOTA) foundation PLM, ProtTrans [18], to generate biologically meaningful embedding for each plasmid-encoded protein as the raw input for our models. Then, we define plasmid sequences as a language using the vocabulary of proteins and leverage the powerful BERT model [19] to capture the genetic structures of plasmids. More specifically, we formulate the GO term prediction as a multi-label token classification task in natural language processing (NLP), where each protein token is assigned one or more GO term labels. Additionally, to increase precision and filter high-confidence predictions, we integrate a self-attention confidence weighting mechanism to learn a confidence score for each predicted GO term. Thus, users can conduct automatic function annotation based on our predictions with associated confidence values. We rigorously evaluated PlasGO on the curated RefSeq dataset and a case study involving two well-studied conjugative plasmids. PlasGO consistently demonstrated superior performance across all experimental results, including the prediction of GO terms for novel proteins. We directly applied PlasGO to 678,197 proteins in the RefSeq plasmid sequences. PlasGO successfully extended high-confidence GO terms for over 95% of the unannotated proteins, which can provide important data for downstream plasmid researches.

## Materials and methods

### Design rationale

Based on the sequential features of protein sequences and the modular characteristics of plasmids, we designed a hierarchical architecture that considers both the local context within a protein sequence and the global context across different proteins. Following the language analogy in NLP, we establish two types of sentences: plasmid sentences (global), in which encoded proteins serve as tokens, and protein sentences (local), where individual AAs act as tokens. Given that a series of powerful PLMs [20][18][21] have demonstrated exceptional performance in extracting biologically meaningful embeddings for protein sentences, our primary contribution lies in the global learning of plasmid sentences. As shown in Figure 2, we frame the GO term prediction for plasmid-encoded proteins as a token classification task in NLP, where GO term labels are assigned to individual protein tokens within a plasmid sentence. Correspondingly, we leverage the BERT model as the central component of PlasGO to capture the structure of plasmid sentences.

**Figure 2.**
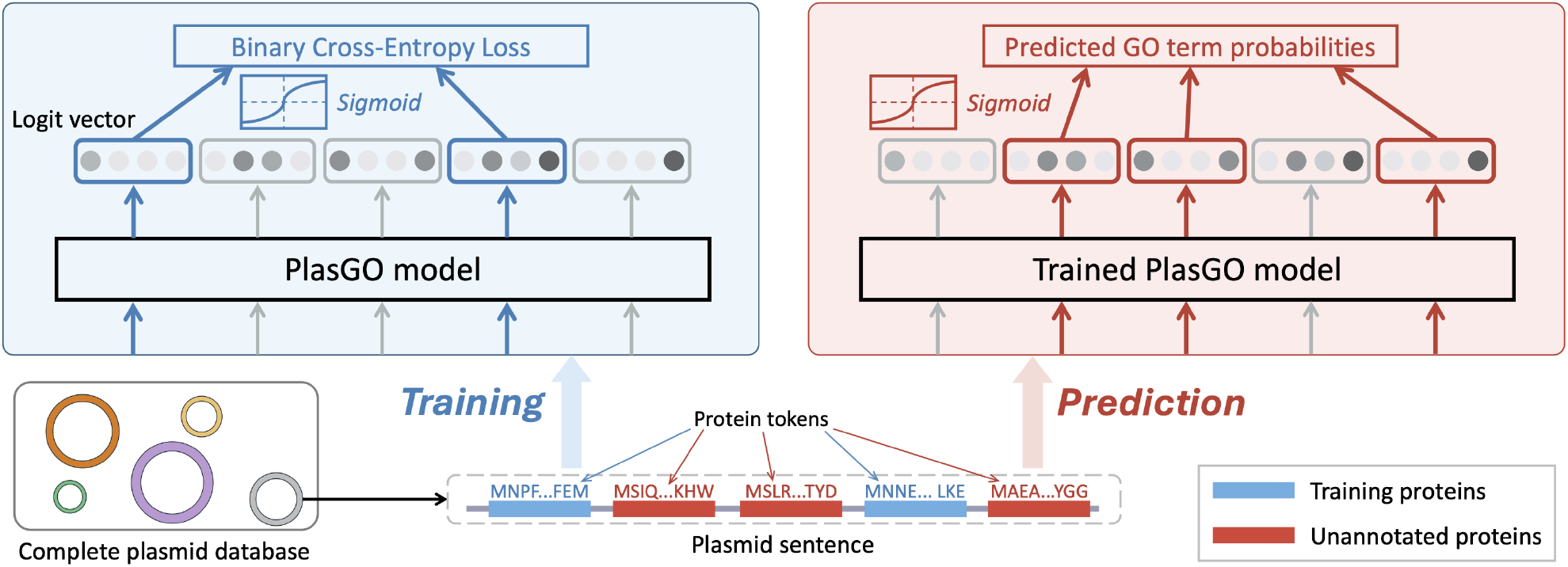
The workflows of PlasGO with a toy plasmid encoding five proteins as input during the training and prediction phases. The figure focuses on the token classification framework, emphasizing the learning of global context across different proteins, which are represented by the per-protein embeddings learned by PLM. Therefore, the toy plasmid is defined as a sentence comprising five protein tokens, and the objective of PlasGO is to predict GO terms for each protein token within the plasmid sentence. Two of the five proteins (depicted in blue) have GO annotations and are included in the training set, while the remaining three (depicted in red) represent proteins without any GO annotation. Throughout both phases, PlasGO learns a vector (passed through sigmoid) for each protein, indicating the probabilities of its corresponding GO terms.

The method design of PlasGO incorporates two distinctive features. First, due to the fact that a protein can be annotated with multiple GO terms, PlasGO is designed as a multi-class, multi-label token classification approach. In other words, the BERT model predicts multiple labels for each protein token, which sets it apart from traditional token classification tasks such as named entity recognition (NER) [22]. Second, PlasGO accepts the same plasmid as input for both supervised training and prediction while focusing on different proteins in each phase. The workflows are depicted in Figure 2 using a toy plasmid. During training, unannotated proteins are masked, and Binary Cross-Entropy (BCE) loss is calculated by comparing the learned GO term probability vectors of the training proteins against their actual GO annotations. In the prediction phase, the trained PlasGO model generates prediction results for all unannotated proteins. In essence, the central concept behind PlasGO is to enhance the function prediction of unannotated plasmid-encoded proteins by leveraging plasmid-level contextual information. On the other hand, even in scenarios where all nearby proteins of a particular protein lack annotations, PlasGO still works by directly predicting GO terms for that protein using the local PLM embeddings.

### Overview of the PlasGO model

As shown in Figure 3, the PlasGO model takes as input a sentence representation of the plasmid, wherein translated proteins (tokens) are arranged in the same order as their encoding in the plasmid. Consequently, the PlasGO model generates high-confidence predicted GO terms for each protein as its output. Specifically, the PlasGO model consists of three modules executed linearly, with the output of each module feeding into the next. The first is the preprocessing module, responsible for generating original per-protein embeddings. This is accomplished by utilizing a pre-trained foundation PLM, which extracts biophysical features of the input protein sequences. The second module is a BERT model. By capturing the contextual information at the plasmid sentence level, the BERT model enhances the original embeddings, encompassing a deeper understanding of the functional characteristics of the proteins. The final is the classifier module, designed explicitly for multi-label token classification. Within this module, we employ a combination of a Fully Connected (FC) layer and a self-attention confidence weighting mechanism. As a result, for each protein token, the classifier generates a GO term probability vector along with a confidence score vector of the same dimension. By removing predictions with low confidence scores, the PlasGO model achieves enhanced accuracy in predicting the GO terms. In the subsequent sections, we will provide a more detailed description of the three modules.

**Figure 3.**
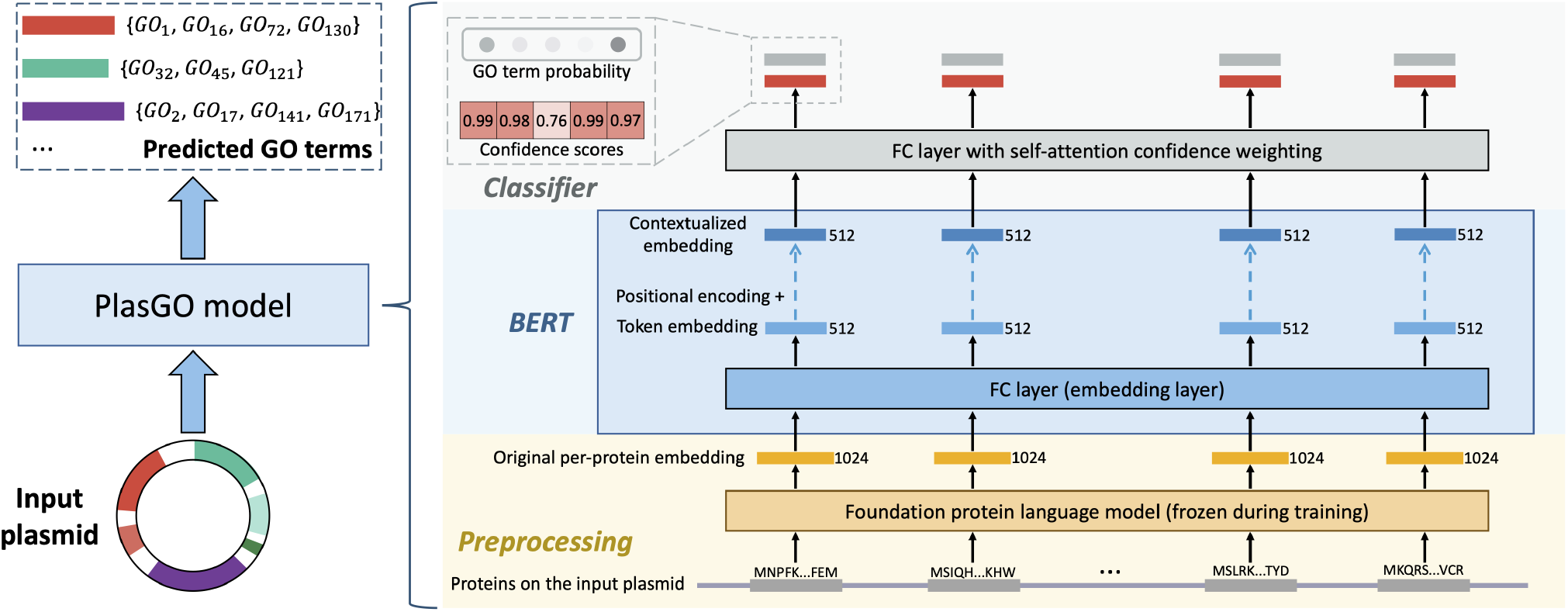
The pipeline of the PlasGO model. The PlasGO model takes as input a series of proteins encoded in a plasmid, and its output comprises high-confidence GO term annotations in nominal format for these proteins. On the right side, the three main modules of the PlasGO model, namely preprocessing, BERT, and classifier, are displayed in a bottom-to-top arrangement. In the preprocessing stage, the proteins (represented as multiple gray bars) are organized in the same order as their encoding in the plasmid. These proteins are then fed into the foundation PLM to extract biologically meaningful embeddings (represented by yellow bars). Next, in the BERT module, an FC layer is utilized to transform the original embeddings learned by PLM into token embeddings (represented by light blue bars). Additionally, multiple Transformer encoders are employed to capture the global context between protein tokens. Lastly, using the learned contextualized embeddings (represented by deep blue bars), the classifier predicts a GO term probability vector (represented by a light gray bar) for each protein token. Simultaneously, a corresponding confidence score vector (represented by a red bar) of the same dimension is also generated. Only the GO terms with both high predicted probabilities and high confidence scores are retained as nominal-format GO term annotations for the respective protein.

### Extract original per-protein embedding with foundation PLM

In this section, we utilize ProtTrans to generate embeddings for each plasmid-encoded protein. Drawing inspiration from concepts in NLP, ProtTrans considers entire proteins as sentences, where individual AAs are analogous to words (tokens). As depicted in Figure 4, the input toy protein sequence “MNPF” is initially tokenized into an array of individual AA tokens [‘M’, ‘N’, ‘P’, ‘F’]. Subsequently, all the AA tokens undergo an embedding layer incorporating positional encoding. This will convert the AA tokens into high-dimensional vectors (1024 dimensions). The resulting embedded vectors are then fed into an *L*-layer Transformer encoder, which captures the semantic meaning of individual AA tokens and their contextual relationships within the protein sequence. In the final step, the output of the last encoder layer for each AA token (per-residue embedding) is concatenated and pooled using a Global Average Pooling (GAP) operation [23]. This involves taking the average of each per-residue embedding, resulting in the final per-protein embedding.

**Figure 4.**
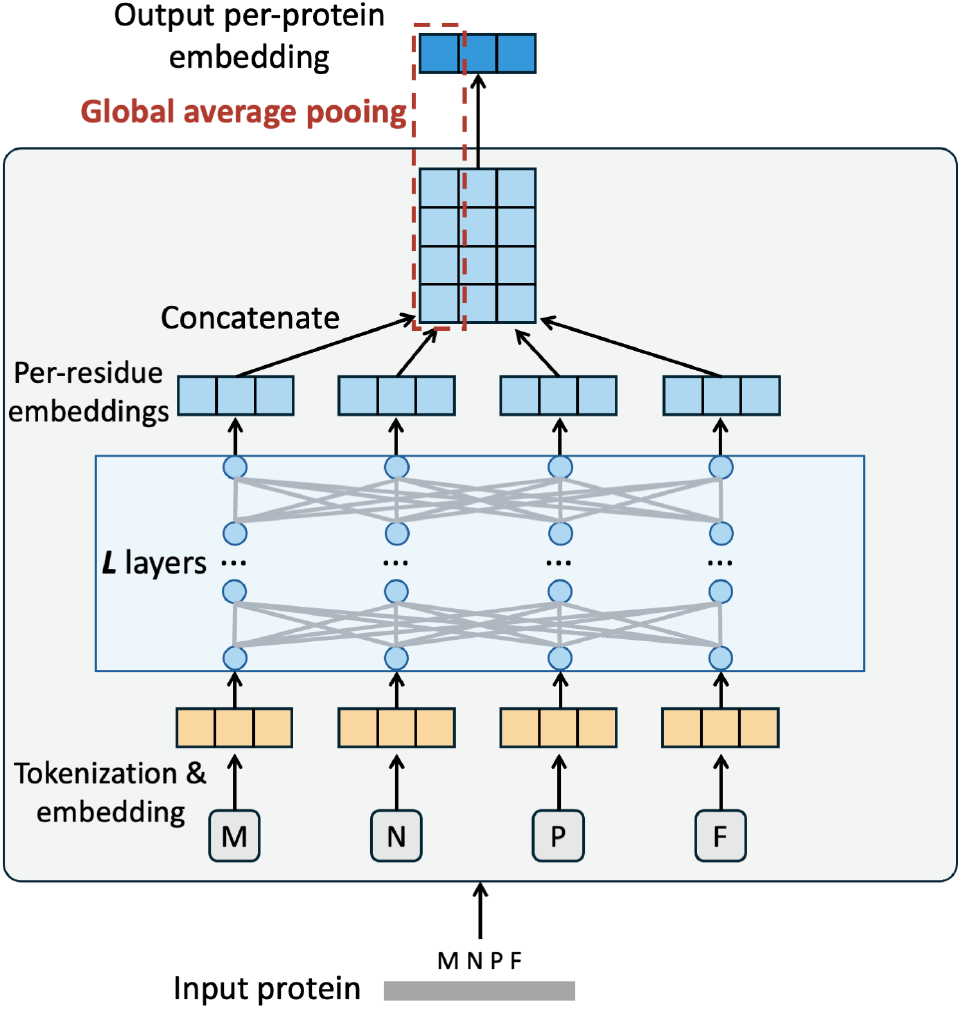
An overview on how the ProtTrans model generates the per-protein embeddings for input plasmid-encoded proteins.

ProtTrans provides several pre-trained models with different parameter numbers and architectures. We selected ProtT5-XL-U50 for PlasGO due to its superior performance across various downstream prediction tasks, as demonstrated in the study by Elnaggar et al. [18]. ProtT5-XL-U50 is built upon the T5-3B model architecture [24] and was trained using the MLM approach [19] on the UniRef50 dataset, which contains 45 million protein sequences. In this work, we employ the ProtTrans model in a feature extraction manner rather than fine-tuning it, which means the per-protein embedding extraction process can be considered as a preprocessing step. Prior to both training and prediction, all the involved proteins are input into ProtTrans, and the resulting per-protein embeddings are saved for later utilization.

### Capture contextual information on plasmids with BERT

We implement our BERT module following the standard BERT architecture [19] with three modifications to accommodate our method design. First, we exclude the two tokens ‘[CLS]’ and ‘[SEP]’ defined in standard BERT, as they are not utilized for token classification. Second, as mentioned earlier, our BERT module is primarily designed to capture functionally related segments in plasmids, similar to how phrases are learned in NLP. Thus, a limited number of layers (Transformer encoders) *L* is adequate for capturing ‘phrase’-level information in plasmid sentences [25]. Specifically, we set *L* = 4 for the MF and BP categories, and *L* = 2 for the CC category, considering the relatively smaller dataset size and label size for CC. Third, we remove the token embedding layer in the standard BERT, which maps token indices to vectors, as we do not represent protein tokens in an indexed form. Instead, in the previous module, we have preprocessed the embedding step using ProtTrans, where an original embedding is extracted for each sequential protein token (AA sequence). Correspondingly, we incorporate an FC layer as the initial layer of our BERT module, which is trained to transform the original embeddings into function-related token embeddings. Next, position embeddings are introduced and combined with the token embeddings:

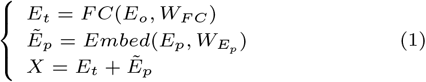

In this work, we set the maximum length of protein tokens in a plasmid sentence to 56, as it represents the median length observed in our curated database of complete plasmids. For sentences with fewer than 56 tokens, the ‘[PAD]’ tokens will be padded at the end of the sentences, and they will be masked during the training. In addition, we utilize a hidden size of *H* = 512 and a number of self-attention heads of *A* = 8 for the Transformer encoders. Therefore, *E*_*o*_ ∈ ℝ^56*×*1024^ is the original embeddings extracted by ProtTrans for the 56 protein tokens, while *E*_*p*_ ∈ ℝ^56*×*1^ is the position index vector. *W*_*FC*_ ∈ ℝ^1024*×*512^ and 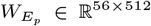 are learnable weight matrices used for the linear transformation of token embeddings and position embeddings, respectively. As a result, both the token embeddings *E*_*t*_ and the position embeddings 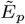 have a dimension of ℝ^56*×*512^, and their sum, denoted as *X*, will be fed into the subsequent Transformer encoders.

### Learn confidence scores for multi-label token classification

In current databases, it is inevitable to encounter incomplete GO annotations because some proteins are annotated with GO terms at varying levels of specificity [26]. Furthermore, accurately predicting certain GO terms poses challenges due to the inherent complexities involved in capturing functional factors such as protein domains or AAs. Consequently, learning a confidence score for each prediction is advantageous as it enables the rejection of uncertain predictions. Inspired by the self-cure network introduced by Wang et al. [27], we integrate a self-attention confidence weighting mechanism tailored for multi-label classification within our classifier module. The detailed architecture of the classifier module is depicted in Figure 5. The contextualized embeddings learned from the BERT module for each protein token will be fed in parallel to the classifier module. Instead of utilizing a single FC layer for multi-label classification, our classifier module incorporates two branches: one for learning logits and another for learning confidence scores. Prior to the final prediction using sigmoid, the logits will undergo attention-weighting based on the learned confidence scores. Intuitively, during training, the model will inherently learn to assign smaller confidence weights to incorrect predictions in order to minimize the loss resulting from these inaccuracies. On the contrary, the true predictions tend to receive larger confidence weights for the logits. The final step involves using both the confidence scores and weighted predicted probabilities to determine the nominal-format GO term predictions, retaining only those predictions characterized by high values in both aspects. Notably, as a modification to the original architecture proposed in [27], we adapt it for the multi-label classification task of GO term prediction by learning a confidence score for each prediction instead of each sample.

**Figure 5.**
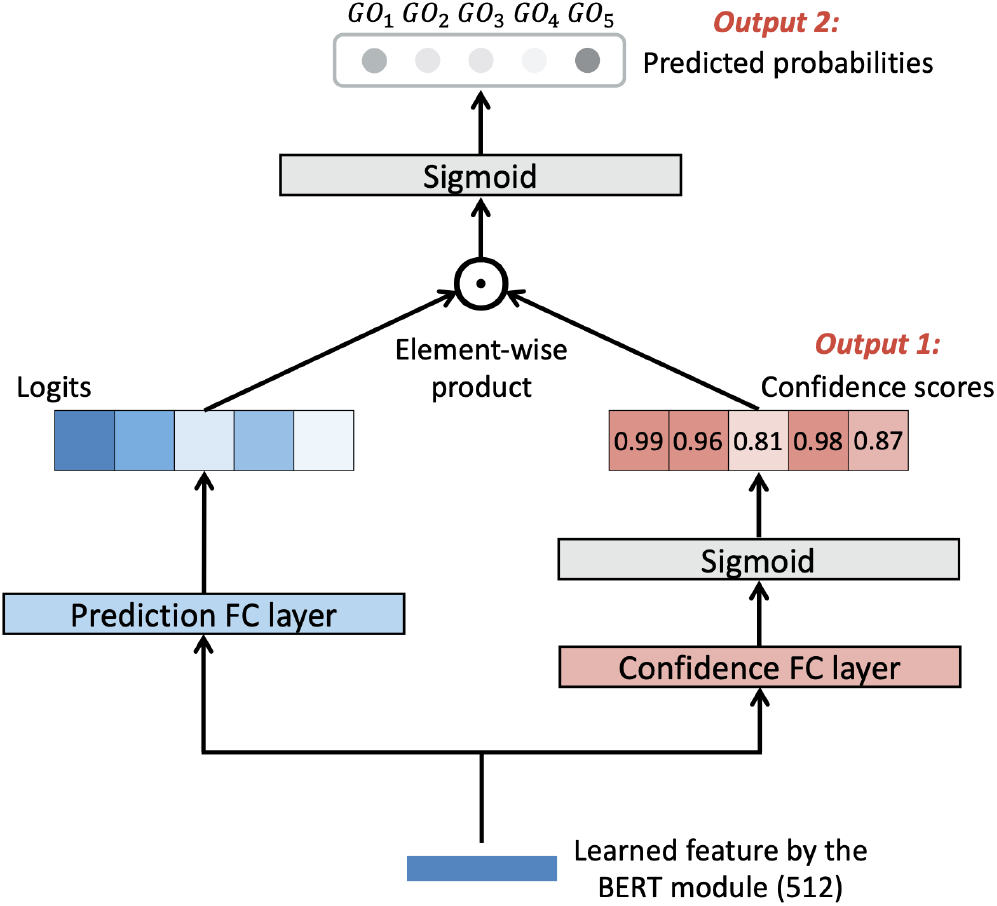
The model architecture of the classifier module incorporating the self-attention confidence weighting mechanism. We utilize a toy label set comprising 5 GO terms for illustration. The figure displays the two branches, with one dedicated to learning logits and the other focused on learning confidence scores, positioned on the left and right sides, respectively. The attention weighting is implemented between the outputs of the two branches to obtain the final multi-label prediction. Both the confidence score vector and the final predicted probability vector will be output by the classifier module to determine the high-confidence GO term predictions in nominal format.

Specifically, the input feature will be forwarded through two FC layers with identical sizes but distinct parameters. These layers consist of a prediction FC layer, responsible for learning logits, and a confidence FC layer, specifically designed to get confidence scores using a sigmoid function. Following that, the confidence scores will serve as the attention weights for the logits through an element-wise dot product:

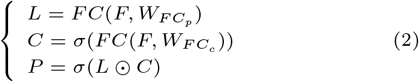

where *F* ∈ ℝ^56*×*512^ is the input features of the 56 protein tokens within one sentence. Here, *n* represents the number of GO term, and *σ* is the sigmoid function. Accordingly, 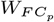 and 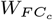 represent the learnable weight matrices, both having dimensions of ℝ^512*×n*^, associated with the prediction FC layer and the confidence FC layer, respectively. Additionally, *L* and *C* denote the learned logits and confidence score matrix, respectively, both having dimensions of R^56*×n*^. The final predicted probability matrix *P* ∈ ℝ^56*×n*^ is obtained by element-wise multiplying (⊙) the matrices *L* and *C*, followed by normalization using the sigmoid function. For the multi-label token classification of PlasGO, we define the logit-weighted binary cross-entropy loss (WBCE Loss) as follows:

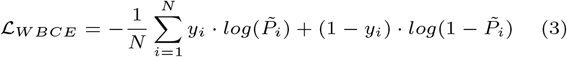

where *N* denotes the total number of predictions, which is equal to 56×*n* for a single sentence. 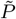 represents the flattened format of the probability matrix *P*, with a dimension of 1 × *N*. *y*_*i*_ represents the true label of the corresponding GO term, where it takes the value 1 if the protein is annotated with that particular GO term, and 0 otherwise.

To further promote the ability of the model to distinguish low-confidence and high-confidence predictions, a rank regularization loss (RR Loss) is incorporated into the total loss function: ℒ_*total*_ = ℒ_*W BCE*_ + ℒ_*RR*_. The calculation of RR Loss and the selection of high-confidence GO term predictions in nominal format are detailed in Supplementary Section S1.

### Data curation and model training

#### RefSeq dataset

We initially downloaded all available plasmids from the NCBI RefSeq database, along with their corresponding protein sequences that were translated from CDSs, with the exclusion of pseudogenes. They can be stored in a dictionary format *plasmid*_*A*_ : [*protein*_*A*1_, *protein*_*A*2_, …], where the keys are plasmids and the values are lists of proteins arranged in the order they are encoded in the respective plasmids. The focus of this work is proteins in regular plasmids. Thus, we only keep plasmids with lengths between 1K and 350K to ensure each plasmid has at least one encoded protein and no megaplasmids are included [28]. The maximum length of protein sequences was restricted to 1K, aligning with other GO term prediction tools such as PFresGO [10]. Besides, the curated proteins’ associated GO terms were also downloaded from the RefSeq database, if available.

Since the GO terms provided in the database are commonly more specific (on the lower levels), we augment the original annotations by propagating all ancestors of the GO terms to their corresponding proteins. As a consequence, the resulting list describing one protein’s function often includes a large number of highly redundant GO terms, which reduces interpretability for users. Several tools have been proposed to tackle this problem by clustering GO terms with high semantic similarity. The measurement of semantic similarity [29] relies on the proximity of two GO terms within the GO DAG. When two GO terms exhibit greater semantic similarity, they tend to be more functionally related. Among these tools, we selected the widely adopted REVIGO [30] to group 1,602 original GO terms into 487 clusters. If at least one GO term within a cluster is annotated to a protein, then that cluster will be assigned to the protein. Importantly, the GO term label clustering strategy is applied to the retraining processes of all benchmarked tools, ensuring a fair comparison of their performance. Afterwards, we deduplicated the proteins by clustering protein sequences using MMseqs2 [31], considering pairs with at least 90% identity and 80% overlap, and only keeping the longest protein from each cluster. In line with the UniRef database [32], the annotations of the kept protein are determined as the intersection of the annotations of all members within the protein cluster. As a result, these two clustering processes have partially corrected misannotations of GO terms in the database.

Consistent with the settings commonly used in GO term prediction tools, we selected the GO term clusters associated with ≥50 proteins (excluding the three root terms) as the labels for our curated dataset. This resulted in 172, 174, and 31 cluster-level labels for the MF, BP, and CC categories, respectively. Subsequently, we devised a data-splitting strategy to simulate the scenario of function prediction for novel proteins. For each GO category, we allocated 10% of the most recently released proteins with GO annotations as the test set. Additionally, we ensure that the novel test set significantly differs from the training set in terms of protein sequences. Therefore, among the remaining 90% annotated proteins, those lacking significant alignments (E-value*>*1e-3) to the test set were assigned to the training set, while others were assigned to the validation set. This splitting strategy poses significant challenges for both PlasGO and other DL methods. The final step involves converting plasmids into sentences for PlasGO’s training and prediction. Plasmids containing more than 56 proteins were divided into multiple segments with an overlap of 14 proteins (1/4 of the maximum length). The curated dataset will be utilized for retraining all benchmarked tools, and a comprehensive overview of its specific details can be found in Table 1.

**Table 1.** The specific information of the curated dataset. In this table, the term ‘sentence’ specifically refers to the plasmid sentence, which is composed of multiple protein tokens. Besides, the ‘# of sentences’ item in the last three columns represents the count of sentences that contain at least one protein from the respective protein set.

| GO category | # of sentences<br>(# of plasmids) | # of deduplicated proteins<br>(# of original proteins) | # of annotated<br>proteins | Training set size<br>(# of sentences) | Validation set size<br>(# of sentences) | Test set size<br>(# of sentences) |
| --- | --- | --- | --- | --- | --- | --- |
| MF | 89,835 (47,871) | 678,197 (1,103,790) | 173,666 | 99,806 (87,198) | 56,491 (44,026) | 17,369 (77,074) |
| BP |  |  | 99,945 | 60,143 (81,068) | 29,768 (72,128) | 10,034 (32,751) |
| CC |  |  | 28,081 | 21,228 (77,877) | 4,045 (18,226) | 2,808 (8,952) |

#### PlasGO model training

We trained three PlasGO models, each specifically designed for one of the three GO categories. The models were trained with a batch size of 32 and a learning rate of 1e-4. Due to the smaller data size and increased susceptibility to overfitting in the CC category, we applied a dropout rate of 0.2 for CC and 0.1 for MF and BP. Additionally, we incorporated a warmup strategy by allocating 5% of the total training steps to gradually increase the learning rate from a small value to 1e-4. Subsequently, the learning rate was linearly decayed to enhance generalization and expedite convergence. Using the NVIDIA GeForce RTX 3090 Blower 24G graphics card, each model underwent 10 epochs of training, with approximate durations of 65 minutes for MF, 59 minutes for BP, and 51 minutes for CC.

## Results

### Experimental setup

#### Metrics

In our benchmark experiments, we assess the performance using two commonly used metrics in the CAFA challenge [33]: the protein-centric *F*_*max*_, which measures the accuracy of assigning GO terms to a protein, and the term-centric area under the Precision-Recall curve (AUPR), which evaluates the accuracy of predicting which proteins are associated with a given GO term. In other words, we first calculate *F*_*max*_ for each protein and AUPR for each GO term separately. Subsequently, we obtain the average of these individual metrics as an overall performance evaluation. Importantly, dataset imbalance can lead to a high *F*_*max*_ but a low AUPR if the tool consistently fails to predict certain low-frequency GO term labels. Therefore, utilizing both metrics ensures a more accurate and comprehensive assessment of the tools’ performance in GO term prediction.

*F*_*max*_ is defined as the highest *F*_1_-score achieved among all probability cutoffs *θ*. To elaborate, we calculate the *F*_1_-score for each *θ* value ranging from 0 to 1, using a stride of 0.01, and select the maximum score as *F*_*max*_:

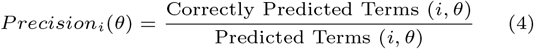

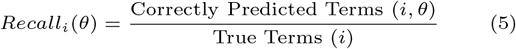

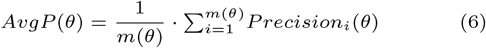

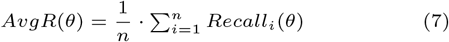

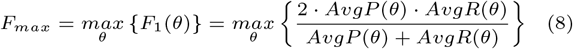

where *Precision*_*i*_(*θ*) and *Recall*_*i*_(*θ*) represent the precision and recall values for protein *i* with the cutoff *θ*. Correspondingly, *AvgP* (*θ*) and *AvgR*(*θ*) represent the average precision and recall values calculated for proteins with at least one predicted GO term (*m*(*θ*)) and all proteins (*n*) respectively. *Predicted Terms* (*i, θ*) denotes the number of GO terms for protein *i* that have a predicted probability greater than *θ. True Terms* (*i*) represents the number of GO terms that are truly annotated for protein *i*.

#### SOTA tools for benchmarking

We compared PlasGO with six deep-learning-based tools that can be used for protein function annotation. These tools encompass DeepGOPlus [8], PFresGO [10], DeepSeq [7], TALE [9], CaLM [17], and an approach trained for accurate protein structure alignments (TM-Vec [34]). Due to our dataset splitting strategy, which resulted in no significant alignment between the test set and training set, we did not include any sequence alignment tools such as Diamond [35] in our comparison. Notably, all six selected SOTA tools can be optimized for GO term prediction for plasmid-encoded proteins using the same curated RefSeq dataset as PlasGO, ensuring a fair and consistent comparison of algorithms. To be specific, we performed model retraining for the first four tools, and in the case of CaLM, we trained a classifier utilizing its learned embeddings. Besides, for TM-Vec, we created a custom database using our curated training set. There are two main reasons for this optimization process. First, some of the labels determined by us do not exist in their default models, making them unable to predict those labels. For example, the default PFresGO can only predict 125 out of 172 (72.67%) MF labels, 141 out of 174 (81.03%) BP labels, and 21 out of 31 (67.74%) CC labels from our label set. Second, the proteins used to train their default models span across various organisms and have less emphasis on plasmids. As a result, their default models exhibit inferior performance compared to our retrained models. For instance, among the subset of labels that can be predicted by the default PFresGO, it achieved *F*_*max*_ scores of 0.6514 and 0.6226 for MF and BP, respectively, while our retrained PFresGO improved significantly to 0.7039 and 0.7189, respectively.

When conducting benchmarking with CaLM, we initially extracted the embeddings for each protein and then trained a Deep Neural Network (DNN) as its GO term prediction classifier. In addition, TM-Vec generates structure alignments in a format similar to TM-align [36], represented as a triad {*q, t, tmscore*(*q, t*)}. Here, *q* denotes the query protein from the test set, *t* represents the target protein from the training set database, and *tmscore*(*q, t*) corresponds to the template modeling score (TM-score) of the alignment. Alignments with a *tmscore*(*q, t*) below 0.5, indicating a low structural similarity [37], are excluded from further analysis. Inspired by the DiamondScore method proposed by Kulmanov et al. [8], we compute the GO term probability vector for TM-Vec using the following formula:

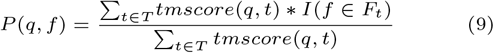

where *P* (*q, f*) is the predicted probability that the query protein *q* is annotated with the GO term *f*. *T* represents the set of target proteins found in the significant alignments of the query protein *q. I*(*f* ∈ *F*_*t*_) is the identity function that returns 1 if the GO term *f* is present in the true annotations *F*_*t*_ of the target protein *t*, and 0 otherwise.

### Performance on the RefSeq test set

We compared PlasGO with the other six tools on our curated RefSeq test set. During the evaluation of PlasGO, if a protein appears in multiple testing sentences, its final probability vector is determined by averaging all its predictions across these sentences. The predicted results of all the tools are shown in Figure 6. Overall, PlasGO attained the highest scores for both *F*_*max*_ and AUPR on all three GO categories. A closer looks show that the top three tools (PlasGO, PFresGO, TM-Vec) all utilize the pre-trained PLM, ProtTrans, to encode protein-level or residue-level features for protein sequences. This observation suggests that the inherent relationships between AAs captured by PLMs can substantially enhance the prediction of GO terms. This aligns with the finding that many proteins execute their functions through spatially aggregated clusters of critical residues [38]. Specifically, PFresGO and TM-Vec utilize residue-level features, while PlasGO leverages protein-level features, resulting in significant time and memory savings, and also scalability to a higher number of proteins. This advantage is achieved by implementing the BERT model to learn from plasmid-level corpora.

**Figure 6.**
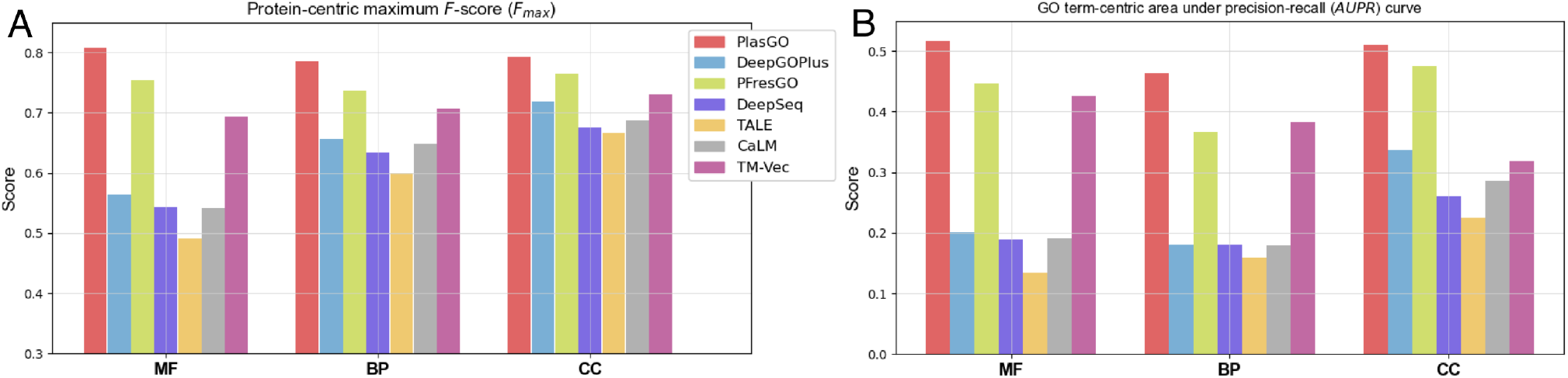
The performance of different tools on the RefSeq test set assessed based on two metrics: A) *F*_*max*_ and B) AUPR, and measured across the three GO categories.

CaLM, despite being a pre-trained PLM, did not surpass the top three tools using ProtTrans, which is consistent with the results reported in CaLM’s paper [17]. Nevertheless, CaLM’s ability to capture proteins’ biochemical characteristics with codons proves particularly advantageous in tasks such as protein abundance prediction. On the other hand, DeepGOPlus achieves the highest performance among the tools trained from scratch. However, it is still challenging for them to compete with the tools trained based on pre-trained PLMs.

### Ablation studies: validating PlasGO’s design rationale

#### Evaluation of BERT module in PlasGO

We first conducted an ablation study to investigate whether the BERT module effectively captures the contextual information on plasmids and improves the GO term prediction. Specifically, we compared the performance of three different classification methods, all utilizing protein embeddings from ProtT5 as input. The first method serves as the baseline, entailing the training of a 3-layer DNN classifier for each GO category using the identical dataset as PlasGO. This method does not leverage any plasmid-level contextual information as it predicts each protein independently. The second method involves using PlasGO for prediction by inputting a single testing protein token. In other words, each testing sentence has a length of 1, with 55 ’[PAD]’ tokens padded at the end of each testing protein token. In this approach, the proteins are also predicted individually. Thus, the main effective component is the embedding layer, while the attention blocks remain inactive. The third method is the standard PlasGO, which fully incorporates our design rationale. The experimental results are shown in Table 2. The standard PlasGO achieved the best performance on all the GO categories, indicating that the contextual information captured by the BERT module actively contributes to improving the GO term prediction. Additionally, the second method has a better performance compared to the DNN classifier. This suggests that despite the inactive attention blocks, the embedding layer still captured partial contextual information.

**Table 2.** The performance of different classification methods using ProtT5 embeddings as input on the RefSeq test set. The first method involves training a 3-layer DNN using ProtT5 embeddings as input, utilizing the same curated RefSeq dataset aligned with PlasGO. The second method deviates from the standard PlasGO approach solely during the prediction phase. Specifically, a single test protein token is treated as a test sentence with a length of 1, which is then inputted into the PlasGO model. Consequently, the attention mechanisms are disabled for the second method.

| Method | GO category | $F_{max}$ | AUPR |
| --- | --- | --- | --- |
| 3-layer DNN classifier | MF | 0.7662 | 0.4550 |
|  | BP | 0.7498 | 0.3903 |
|  | CC | 0.7778 | 0.4639 |
| PlasGO (single testing token) | MF | 0.7808 | 0.4950 |
|  | BP | 0.7512 | 0.4088 |
|  | CC | 0.7992 | 0.4406 |
| PlasGO (standard) | MF | 0.8070 | 0.5165 |
|  | BP | 0.7855 | 0.4638 |
|  | CC | 0.7926 | 0.5109 |

#### Evaluation of the foundation PLMs

Given that the protein embeddings extracted by the foundation PLM are the sole input to PlasGO’s BERT module, it is critical to ensure that the input embeddings are highly informative. To this end, we evaluated the performance of PlasGO by utilizing embeddings extracted from four distinct ProtTrans models, which were constructed based on four prominent language models in NLP: T5 [24], BERT [19], XLNet [39], and Albert [40]. The results align closely with the per-protein prediction tasks reported in ProtTrans’s paper [18] (Table 3). PlasGO achieved the best results using ProtT5, while the differences in performance between PlasGO using other PLMs were minimal. This suggests that the model size of ProtT5 (3B) is optimal for learning the most informative embeddings from the vast number of training proteins (e.g., 45M in the UniRef50 database). Thus, we select ProtT5 to extract the original per-protein embeddings for PlasGO.

**Table 3.** The performance of PlasGO trained with different foundation PLMs on the RefSeq test set.

| Pre-trained PLM | GO category | $F_{max}$ | AUPR | Parameters |
| --- | --- | --- | --- | --- |
| ProtT5 | MF | 0.8070 | 0.5165 | 3B |
|  | BP | 0.7855 | 0.4638 |  |
|  | CC | 0.7926 | 0.5109 |  |
| ProtBert | MF | 0.7462 | 0.4532 | 420M |
|  | BP | 0.7447 | 0.3813 |  |
|  | CC | 0.7444 | 0.3915 |  |
| ProtXLNet | MF | 0.7396 | 0.4693 | 409M |
|  | BP | 0.7526 | 0.3622 |  |
|  | CC | 0.7673 | 0.4567 |  |
| ProtAlbert | MF | 0.7541 | 0.4220 | 224M |
|  | BP | 0.7396 | 0.3503 |  |
|  | CC | 0.7679 | 0.4393 |  |

### Visualization of the PlasGO embeddings

A visual comparison between the original embeddings generated by ProtTrans and the contextualized embeddings learned by the BERT module of PlasGO can provide valuable insights into how the design of the BERT module contributes to enhancing GO term prediction. Given that the multi-label GO term prediction can be considered as many individual binary classifications, we conducted visualization experiments focusing on several GO terms that are highly related to plasmid-specific functions. Specifically, we first selected a few representative plasmid-specific proteins, encompassing both core proteins and accessory proteins. Subsequently, we obtained the corresponding GO annotations for these proteins from the Swiss-Prot database. On one hand, we curated a subset of well-studied plasmid core proteins (Supplementary Section S2) from the categories of replication, partitioning, conjugative DNA transfer, exclusion, and type IV secretion system (T4SS), based on the list provided by Thomas et al. [41]. On the other hand, we retrieved high-quality proteins labeled as ’plasmid’ (Supplementary Section S3) from the Swiss-Prot database by utilizing four crucial plasmid accessory functions as keywords: AMR, resistance to heavy metals, new metabolic process, and virulence factors. In this section, we validated the effectiveness of PlasGO in capturing the underlying features for classifying plasmid-specific functions by selecting four GO terms from the two curated lists for each GO category.

For each selected GO term, we categorized the proteins associated with it as positive samples, while considering the remaining proteins not associated with it as negative samples. An inspection of our curated dataset revealed an imbalanced distribution of data across all twelve selected GO terms, with the majority of proteins classified as negative samples. To enhance clarity in the visualization, an equal number of negative samples were randomly sampled to match the number of positive samples for each GO term. We employed t-SNE (t-distributed stochastic neighbor embedding) [42] for 2D visualization of both the original embedding from ProtTrans and the embeddings generated by PlasGO’s BERT module. The comparison of embeddings for the four MF binary classifications is presented in Figure 7, while the comparisons for BP and CC are shown in Supplementary Section S4. To quantitatively evaluate the separation of embeddings between the two classes, we consider the positive proteins and negative proteins as separate clusters. Then, we calculate the silhouette score (ranging from -1 to 1) based on these two clusters. We can observe that the PlasGO embeddings exhibit a clearer separation of protein functions compared to the ambiguous ProtTrans embeddings. This suggests that PlasGO effectively captures the latent features associated with plasmid-specific functions.

**Figure 7.**
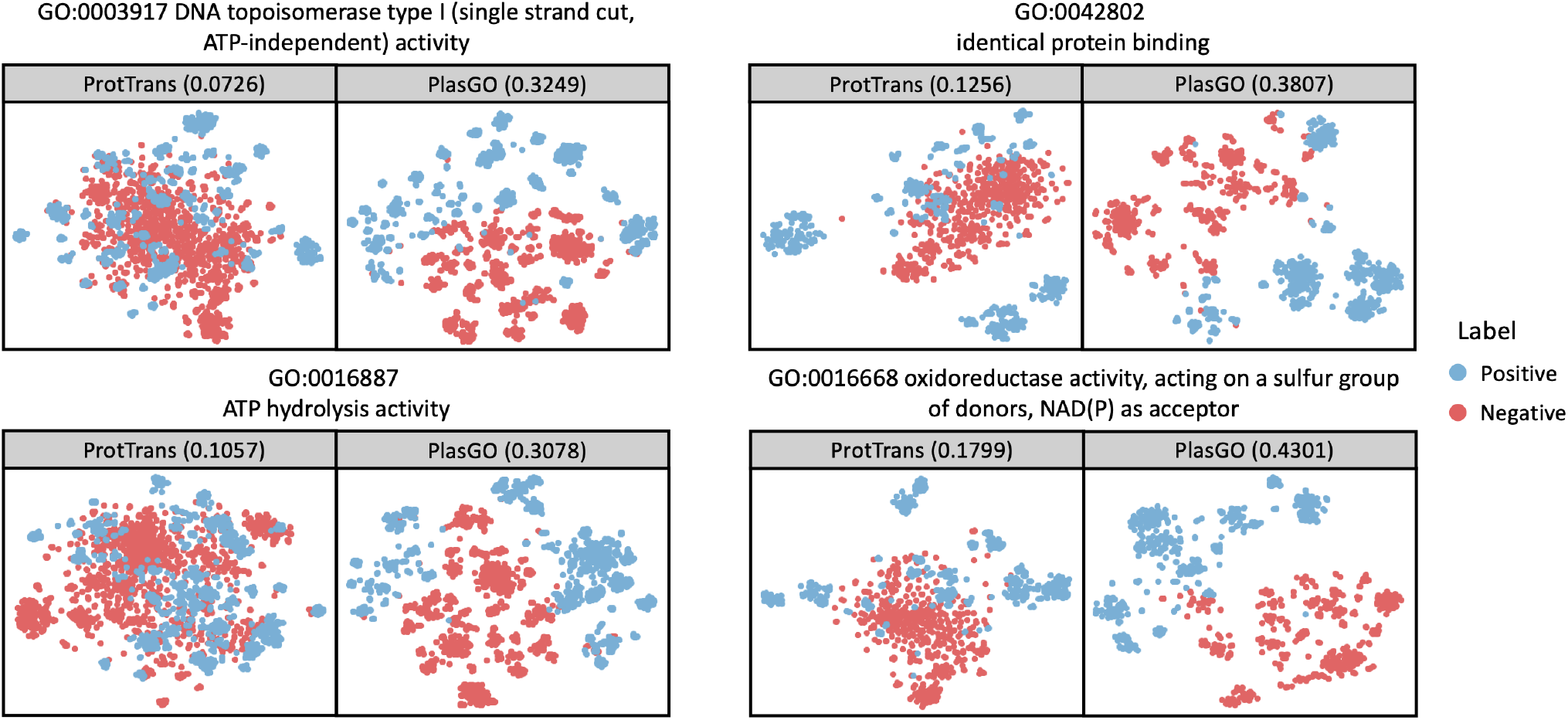
The visualization of protein embeddings generated by ProtTrans and PlasGO for four GO terms. For each GO term, the proteins associated with it and not associated with it are defined as positive and negative samples, respectively. Blue dots: positive samples; red dots: negative samples. The X-axis and Y-axis represent the two dimensions of the embeddings reduced using t-SNE. Additionally, the silhouette score is displayed within brackets on each box.

### Identification of elusive GO term labels

Predicting all GO terms accurately poses a significant challenge due to its complex multi-label classification. To make our tool more practically useful, we prefer to sacrifice prediction resolution for higher precision. Based on the empirical experiments, we identified a few labels that are extremely difficult to predict. Specifically, if the term-centric AUPR score on the validation set of a label falls below 0.3, then this label is considered an elusive label. As a result, a total of 16 out of 172 MF labels, 21 out of 174 BP labels, and 3 out of 31 CC labels have been identified as elusive labels. The detailed list of these elusive labels, along with their corresponding AUPR scores on the test set obtained from the top four tools (PlasGO, PFresGO, TM-Vec, and DeepGOPlus), is provided in Supplementary Table S1. It is notable that the majority of the elusive labels, with the exception of 5 of them (where PlasGO still outperforms the other tools), consistently achieve AUPR scores below 0.3 on the test set across all the top four tools. In summary, this suggests that current data cannot support reliable prediction of these labels.

There are three main reasons why elusive labels are challenging to predict accurately. First, the distribution of GO terms in the training set is imbalanced. As depicted in Supplementary Figure S3, the majority of elusive labels are rare classes, resulting in difficulties for the model to effectively learn their associated patterns. Second, due to our dataset-splitting strategy, the dissimilarity in protein sequences between the training set and test set is considerable (Supplementary Figure S4). As a result, limited features can be used for predicting the novel functions of proteins in the test set. As shown in Supplementary Table S2, we filtered the results of the PlasGO module ablation study (Table 2) to specifically consider the elusive labels. The poor performance of the DNN classifier suggests that even the powerful PLM failed to capture the features necessary for predicting the elusive labels. Third, the identified elusive labels have no overlap with the important plasmid-specific GO terms associated with both plasmid core proteins and accessory proteins, as detailed in Supplementary Sections S2 and S3. Hence, the functions represented by these elusive labels may not adhere to a distinct modularization pattern on plasmids, and as a result, PlasGO was unable to improve the prediction of these elusive labels by leveraging contextual information (Supplementary Table S2).

Since the elusive labels are not the most informative GO terms related to plasmid functions, they will be removed from the results presented in the following sections and prediction outputs for users. However, the majority of these elusive labels have their ancestor terms reversed, as shown in the example DAG structure in Supplementary Figure S5. Additionally, as illustrated in Supplementary Figure S6, the nearest ancestor terms of most elusive labels maintain good performance. This suggests that the removal of the elusive labels achieved a significant increase in overall performance despite sacrificing some prediction resolution.

### Labels of different frequencies and confidence scores

We first conducted a more detailed evaluation of the performance comparisons on the RefSeq test set among the top four tools. Specifically, we focused on a few representative labels, namely the first 10 and last 20 most frequent MF labels in the training set. In general, labels that occur more frequently tend to exhibit better performance. This can be attributed to the abundance of training samples, which allows the models to acquire richer features for effectively distinguishing these labels. This pattern aligns with the results presented in Figure 8, which illustrates that the performance of the top 10 labels is superior and more stable compared to the performance of the last 20 labels. Despite obtaining low AUPR scores for several low-frequency labels, PlasGO consistently outperforms the other three tools across the majority of labels.

**Figure 8.**
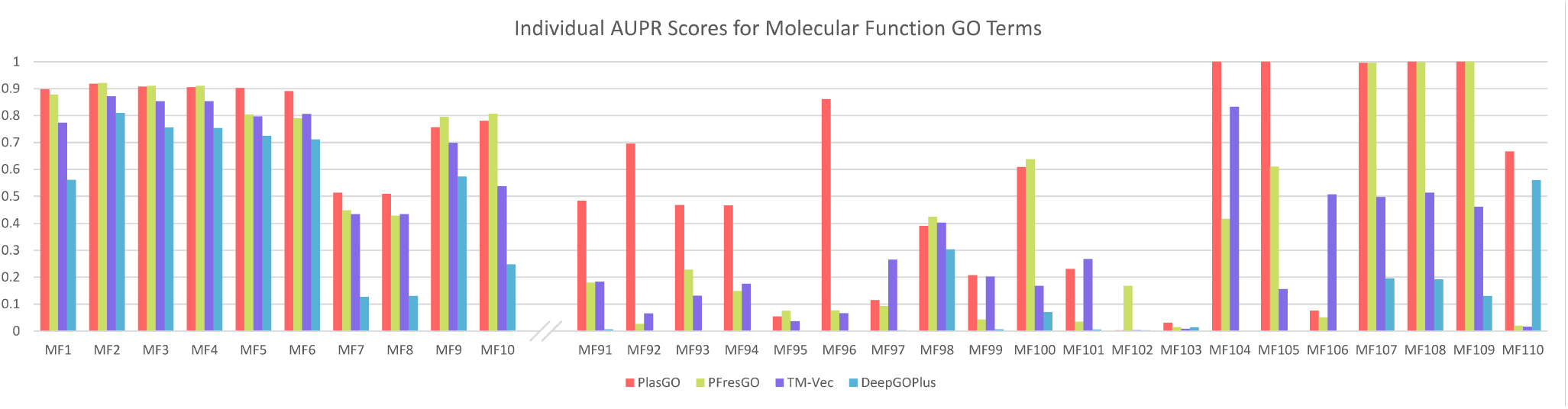
The AUPR comparisons of the top four tools on the first 10 and last 20 MF labels sorted by occurrence frequency in the training set. The performance of all tools fluctuates significantly on low-frequency labels.

Next, we assessed the effectiveness of the learned confidence scores. As described in the Methods section, as the default setting, any prediction with a predicted probability below 0.425 and a confidence score lower than 0.95 will be excluded when calculating the AUPR for the high-confidence mode of PlasGO. To quantify this exclusion, we introduced a new metric called the “prediction rate”, which is calculated by dividing the number of reserved testing proteins by the total number of testing proteins. Figure 9 presents the performance comparison between the original PlasGO and its high-confidence mode for a subset of the MF labels, where the prediction rates are not equal to 1 in the high-confidence mode. We can observe that the high-confidence mode of PlasGO attains higher AUPR scores for the majority of the displayed labels by sacrificing a certain degree of prediction rate.

**Figure 9.**
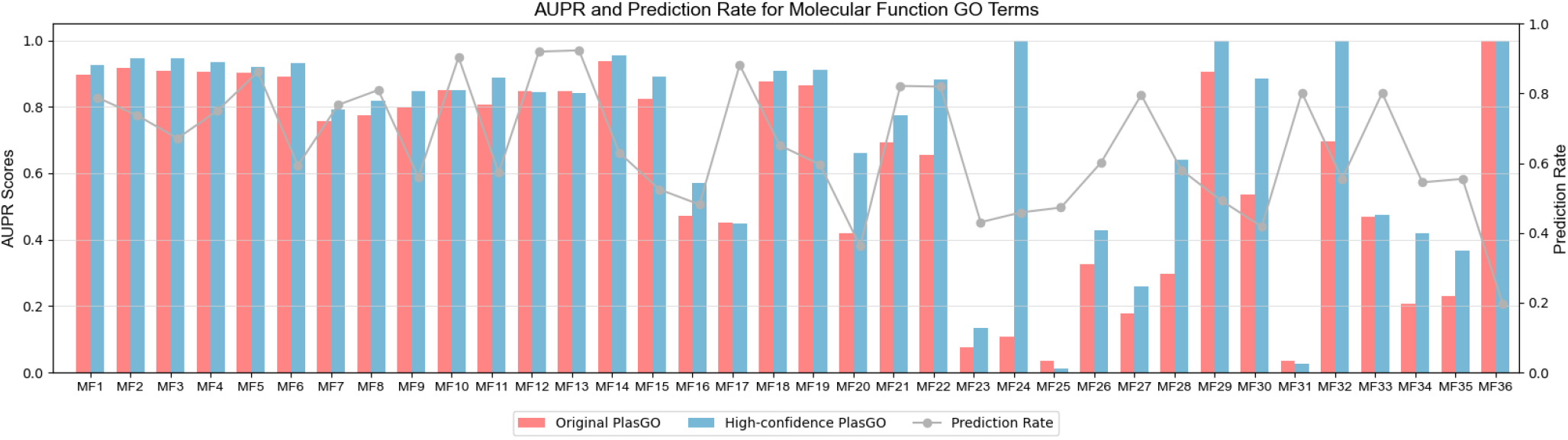
The AUPR comparisons on the MF category between the original PlasGO and the high-confidence mode of PlasGO. In the high-confidence mode, PlasGO filters out some predictions with low learned confidence scores, resulting in higher AUPR scores but lower prediction rates. The gray line shows the prediction rate for the “high-confidence PlasGO” mode. The prediction rates for the original PlasGO are all equal to 1 and thus are not shown in this figure. The corresponding comparison results on the BP and CC categories are shown in Supplementary Section S6.

### Application: automatic GO prediction for unannotated plasmid-encoded proteins in RefSeq

One primary contribution of PlasGO is its utilization of trained models to predict high-confidence GO term annotations for unannotated plasmid-encoded proteins that are not included in the training, validation, and test sets. As shown in Table 4, a large proportion of proteins curated from the RefSeq database (678,197) lack GO term annotations, with percentages of 74.39% for MF, 85.26% for BP, and 95.86% for CC. Hence, we employed PlasGO to predict nominal-format GO term annotations for these unannotated proteins. Consequently, we have successfully assigned high-confidence GO term annotations to the vast majority (all exceeding 95%) of the unannotated proteins. The distributions of the number of newly assigned GO terms for the three GO categories are presented in Supplementary Section S7. Lastly, the precision of the nominal-format GO term annotations was measured on the RefSeq test set, resulting in values of 0.8229, 0.7941, and 0.887 for the MF, BP, and CC categories, respectively. This confirms the high reliability of the newly added GO terms by PlasGO.

**Table 4.**
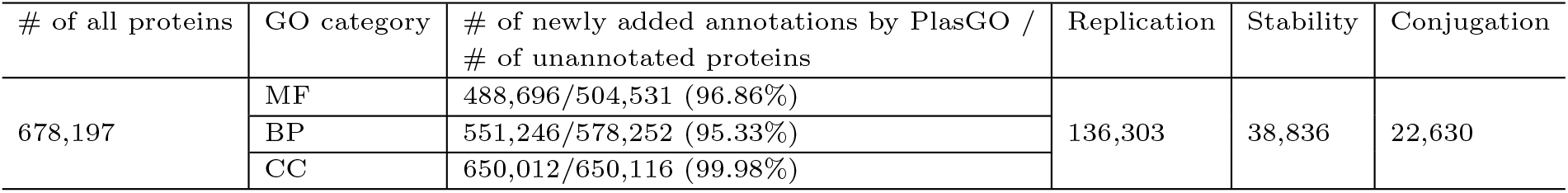
The percentages of newly added high-confidence annotations by PlasGO across three GO categories. The last three columns display the count of proteins that have been newly predicted as having functions related to replication, stability, and conjugation, respectively, utilizing the significant GO term indicators.

| # of all proteins | GO category | # of newly added annotations by PlasGO /<br># of unannotated proteins | Replication | Stability | Conjugation |
| --- | --- | --- | --- | --- | --- |
| 678,197 | MF | 488,696/504,531 (96.86%) | 136,303 | 38,836 | 22,630 |
|  | BP | 551,246/578,252 (95.33%) |  |  |  |
|  | CC | 650,012/650,116 (99.98%) |  |  |  |

To conduct a more detailed analysis of the newly annotated proteins, we additionally provided the count of proteins in Table 4 that can be confidently classified into one of the three main core functions (replication, stability, and conjugation) using PlasGO. In order to achieve this classification, we manually identified the GO terms that exhibit strong relevance as indicators for the three core functions. The comprehensive list of these indicators can be found in Supplementary Section S8. Consequently, any protein predicted to have at least one of these GO term indicators will be classified into the corresponding core function. The results show that PlasGO successfully predicts a significant number of unannotated proteins as plasmid core proteins. Finally, the predicted GO terms for plasmid-encoded proteins were saved in a database alongside PlasGO. Users can first attempt sequence alignment against the database with high identity and coverage cutoffs to directly obtain annotated GO terms. If there is no significant alignment, users can then apply the PlasGO models for GO term prediction.

### Case study: annotations for two well-studied plasmids

We then evaluated PlasGO’s performance in protein function prediction for the two well-studied conjugative plasmids associated with AMR, namely pSK41 from *Staphylococcus aureus* [43] and pOLA52 from *Escherichia coli* [15]. The two complete plasmids, along with their corresponding encoded proteins, were downloaded from the NCBI database using the sequence IDs *NC 005024* and *NC 010378*, respectively. For both plasmids, over half of the encoded proteins possess informative ’gene product’ annotations, excluding those labeled as ’hypothetical protein’ and ’domain-containing protein’. However, only a very small portion of these proteins (4 out of 76) have GO annotations available. Therefore, the ’gene product’ annotations can offer a potential scope of the GO term ground truth, which aids in evaluating the prediction precision of PlasGO for the unannotated proteins. The detailed information on the proteins, including their protein IDs and ’gene product’ annotations, can be found in Supplementary Section S9, presented in the order corresponding to their encoding in the two plasmids. As shown in Figure 10, to obtain the potential ground truth scope, we first retrieved the relevant proteins from the UniProtKB database by using the ’gene product’ annotation as the keyword phrase (e.g., ’MobA/MobL family protein’). Following that, the collected GO terms associated with these retrieved proteins were considered as the potential ground truth set for the corresponding protein. We can then estimate the prediction precision of PlasGO for each protein by calculating the ratio of the number of predicted GO terms in the potential ground truth to the total number of predicted GO terms.

**Figure 10.**
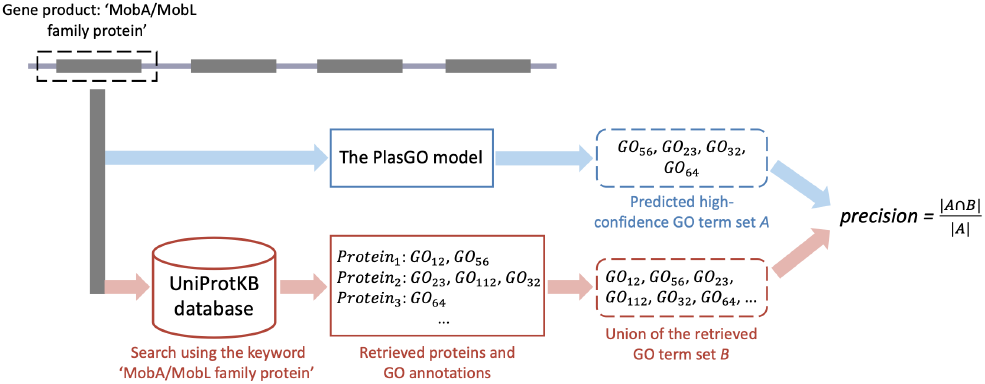
The pipeline of conducting the case study experiment. Only the proteins annotated with informative ’gene product’ annotations were considered in this experiment. In this figure, we present an example evaluation of a protein annotated as ’MobA/MobL family protein’. The blue path illustrates the prediction process of PlasGO, wherein a set of high-confidence GO terms *A* is generated by our algorithm. The red path below depicts the process of obtaining the potential scope of the GO term ground truth. Specifically, we conducted a search in the UniProtKB database using the phrase ’MobA/MobL family protein’ as the keyword. Subsequently, the proteins that matched the keyword, along with their corresponding GO annotations, were retrieved. Finally, we defined the union of the retrieved GO terms as the potential ground truth set *B*. The prediction precision of PlasGO is determined by calculating the ratio of the number of proteins in set *A* that are also present in set *B* to the total number of predicted high-confidence proteins (|*A*|).

Figure 11 illustrates the number comparison between the predicted GO terms (merged from the MF, BP, and CC categories) by PlasGO and the original GO terms available in the RefSeq database, along with the corresponding estimated precision. PlasGO successfully added predicted GO terms to all the proteins and demonstrated a promising estimated precision for the majority of them. It is noteworthy that the increase in the number of GO terms for the four originally annotated proteins can be attributed to PlasGO assigning GO terms to previously unrepresented categories. For example, the 31st protein (*WP 012291478*) in plasmid pOLA52 had only 9 GO term labels in the MF category initially. PlasGO predicts an additional 4 BP terms and 4 CC terms for this protein, resulting in a total of 17 annotated GO terms.

**Figure 11.**
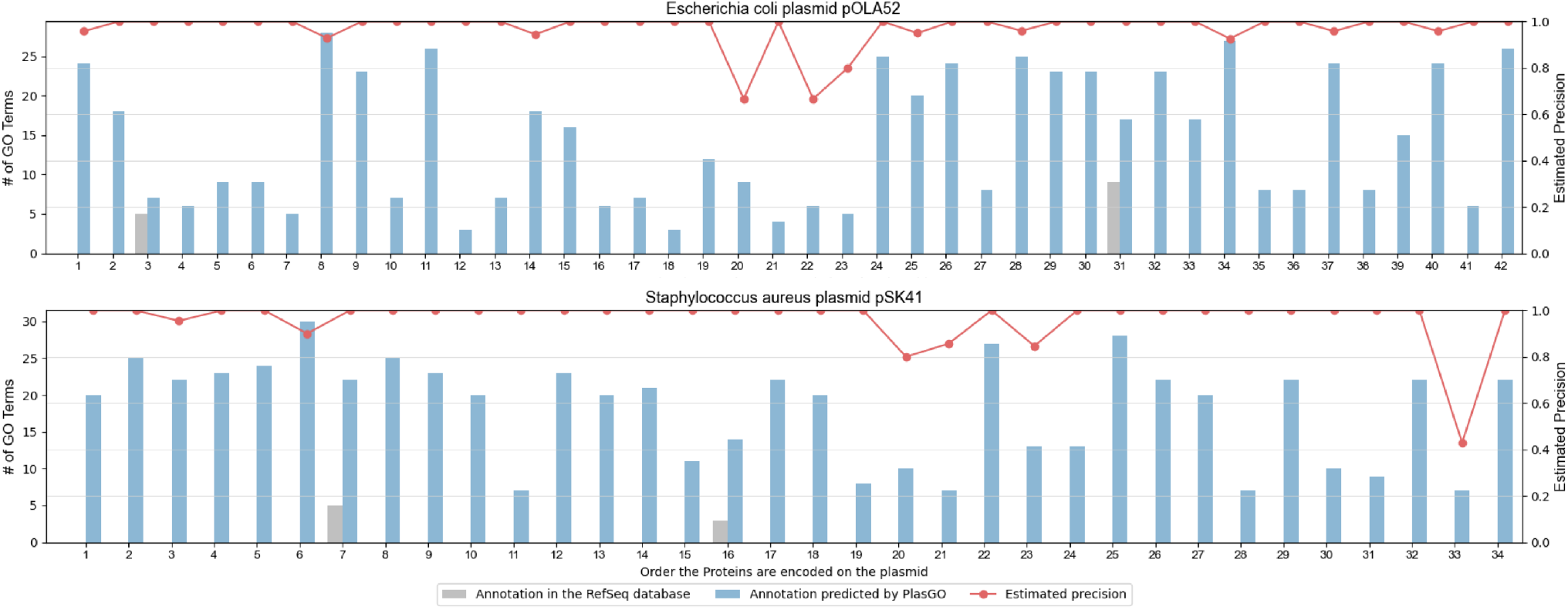
The comparisons between the number of GO terms predicted by PlasGO for the proteins encoded in the two well-studied plasmids and the number of original GO terms available in the NCBI RefSeq database. The indices on the x-axis represent the order in which proteins are encoded in the plasmids. The red lines in the figure represent the precision estimated for each protein using the potential ground truth set retrieved from the UniProtKB database.

## Discussion

In this study, we presented a tool named PlasGO aiming to provide GO term-based functional annotation for largely uncharacterized plasmid-encoded proteins. Due to segment transfer facilitated by recombination events or the movement of MGEs, plasmids frequently demonstrate a modularization pattern, wherein proteins with similar functions tend to be positioned in close proximity to each other. To leverage this characteristic, we represented each plasmid as a sentence and functionally related segments as phrases, both composed of protein tokens. We then formulated the GO term prediction for plasmid-encoded proteins as a multi-label token classification task, utilizing the BERT model. PlasGO is specifically designed with a hierarchical architecture. It utilizes a foundation PLM to capture the local context within a protein sequence while employing a BERT model to capture the global context across different proteins. Furthermore, a classifier module featuring a self-attention confidence weighting mechanism is incorporated to generate confidence scores during prediction, thereby ensuring more reliable results.

We rigorously evaluated PlasGO using a series of experiments and benchmarked its performance with other SOTA tools. Specifically, to mimic the real-world challenges in GO term prediction for novel proteins that cannot be characterized with homology search due to low sequence identity, we constructed a test set that lack significant sequence alignments to the training set. In order to maintain fairness in the comparisons, we retrained models or classifiers for all the other tools by using the same training set as PlasGO. The results from all the experiments consistently demonstrate the superior performance of PlasGO compared with other SOTA tools. Besides, we conducted a careful ablation study to investigate the contribution of different components of PlasGO to performance improvement. These prove that the advantage of PlasGO is attributed to its algorithm design rather than the mere augmentation of training data. Furthermore, the benchmark results indicate that PlasGO has the potential to be extended beyond plasmid-specific applications and applied to general GO term prediction tasks. In the final phase, we utilized PlasGO to predict high-confidence GO terms for the complete set of available plasmid-encoded proteins, totaling 678,197 proteins. Remarkably, over 95% of previously unannotated proteins were successfully assigned new GO terms across all three GO categories. The predicted high-confidence GO terms for all plasmid-encoded proteins will be compiled into a database, which serves as a contribution to the community interested in plasmids.

Despite the notable improvement in function prediction, PlasGO does have two limitations. First, only 29.34% of the proteins in the current database possess GO term annotations. In the most extreme cases, only a single protein token in a sentence may have training labels. This limitation restricts the model’s ability to effectively learn plasmid patterns and hampers their generalization capabilities. Second, despite the high quality of the GO annotations, different proteins may vary in their level of annotation. In other words, some proteins may have annotations at a very detailed level, while others may be annotated to broader, higher-level GO terms. Consequently, the multi-label classification task in PlasGO is inevitably affected by the issue of missing labels, leading to the introduction of label noise. The relabeling method presents a potential approach to address both limitations. Specifically, after the model has undergone training for a specified number of epochs or has converged, labels with high predicted probabilities and confidence scores can be assigned as the ground truth for their corresponding proteins, regardless of whether these proteins were initially annotated or unannotated. Consequently, as training progresses, an increasing number of protein tokens will no longer be masked and will actively contribute to the loss computation. On the other hand, the model will be fully utilized to rectify the missing labels within the dataset, enhancing the accuracy of the predictions.

The architecture of PlasGO is flexible, enabling it to adapt to various functional annotation schemes for plasmid-encoded proteins. On one hand, instead of using GO terms, alternative training labels such as Enzyme Commission (EC) numbers [44], UniProtKB keywords [45], or even customized plasmid-specific protein function classes (e.g., replication and conjugation, though manual labeling efforts may be required) can be employed. On the other hand, PlasGO holds the potential to enhance the performance of the three principal plasmid typing schemes by predicting the classes of the associated proteins [5]. Specifically, Rep typing is based on replication initiation proteins, MOB typing relies on relaxases, and MPF typing is centered around T4SS proteins.

## Data Availability

PlasGO is implemented in Python, which can be downloaded at both figshare (https://figshare.com/articles/journal_contribution/Source_codes_for_PlasGO/26123686) and GitHub (https://github.com/Orin-beep/PlasGO). The curated RefSeq dataset, utilized for training the PlasGO models and benchmarking, as well as the high-confidence GO database for plasmid proteins generated by PlasGO, are available for access on Zenodo (https://zenodo.org/records/12542525).

