## Supplementary data for "PlasGO: enhancing GO-based function prediction for plasmid-encoded proteins based on genetic structure"

June 2024

#### 1 Supplementary Methods: calculation of RR Loss and selection of nominal-format GO term predictions

To compute $\mathcal{L}_{RR}$, the confidence scores output in a batch are initially sorted in descending order. They are then divided into two groups, with a ratio of 70% for the high-confidence group and 30% for the low-confidence group. $\mathcal{L}_{RR}$ is designed to optimize the difference between the mean of the high-confidence group ($\mu_h$) and the mean of the low-confidence group ($\mu_l$), with the goal of approaching a specified hyperparameter $\delta_1$ ($\delta_1 = 0.15$ by default):

$$\mathcal{L}_{RR} = max\left\{0, \delta_1 - (\mu_h - \mu_l)\right\} \tag{1}$$

Importantly, alongside the predicted probabilities $P$, the classifier module also outputs the confidence scores $C$ to assist in determining the GO term predictions in the format of nominal data, such as $protein_A : [GO_i, GO_j, GO_k]$. To illustrate, we demonstrate the determination of $protein_A$'s annotation with GO term $i$ using the predicted probability $P_{Ai}$ and confidence score $C_{Ai}$. Since positive proteins are infrequent for most GO terms, the trained model tends to be conservative in predicting positives. Consequently, if $P_{Ai}$ exceeds a predefined cutoff $\delta_2$ ($\delta_2 = 0.425$ by default), the prediction is considered confident enough, irrespective of the value of $C_{Ai}$, and GO term $i$ is assigned directly to $protein_A$. Alternatively, GO term $i$ is assigned to $protein_A$ only when both conditions, namely $P_{Ai} > \delta_3$ and $C_{Ai} > \delta_4$, are satisfied ($\delta_3 = 0.3$ and $\delta_4 = 0.95$ by default).

### 2 GO term annotations of representative plasmid core proteins

| Function | Entry name | Protein name | GO terms (Molecular Function) | GO terms (Biological Process) | GO terms (Cellular Component) |
|---|---|---|---|---|---|
| Replication | TRFA_ECOLX | Plasmid replication initiator protein TrfA | GO:0003677 DNA binding | GO:0006260 DNA replication<br>GO:0006276 plasmid maintenance | GO:0005886 plasma membrane |
| | SSBF_ECOLI | Single-stranded DNA-binding protein | GO:0003697 single-stranded DNA binding | GO:0006260 DNA replication | |
| Partitioning | PARM_ECOLX | Plasmid segregation protein ParM | GO:0042802 identical protein binding | GO:0030541 plasmid partitioning | |
| | PARB4_ECOLX | Protein ParB | GO:0003677 DNA binding<br>GO:0004519 endonuclease activity<br>GO:0004527 exonuclease activity | GO:0030541 plasmid partitioning | GO:0005576 extracellular region |
| Conjugative DNA transfer | TRAI1_ECOLI | Multifunctional conjugation protein TraI | GO:0003677 DNA binding<br>GO:0003678 DNA helicase activity<br>GO:0003917 DNA topoisomerase type I (single strand cut, ATP-independent) activity<br>GO:0005524 ATP binding<br>GO:0016887 ATP hydrolysis activity<br>GO:0046872 metal ion binding | GO:0008152 metabolic process | GO:0005737 cytoplasm |
| | TRAC5_ECOLX | DNA primase TraC | GO:0003697 single-stranded DNA binding<br>GO:0016779 nucleotidyltransferase activity | GO:0006260 DNA replication | GO:0000428 DNA-directed RNA polymerase complex |
| | TRAD1_ECOLI | Coupling protein TraD | GO:0005524 ATP binding | GO:0009291 unidirectional conjugation | GO:0005886 plasma membrane |
| Exclusion | TRAT1_ECOLI | TraT complement resistance protein | | | GO:0009279 cell outer membrane |
| | TRAS1_ECOLI | Protein TraS | | | GO:0005886 plasma membrane |
| Type IV secretion system | PIL1_ECOLI | Pilin | | | GO:0005576 extracellular region<br>GO:0005886 plasma membrane |
| | TRAL1_ECOLI | Protein TraL | | GO:0009297 pilus assembly | GO:0009279 cell outer membrane |
| | TRBE_RHIRD | Conjugal transfer protein TrbE | GO:0005524 ATP binding<br>GO:0016887 ATP hydrolysis activity | | |
| | TRBL_RHIRD | Conjugal transfer protein TrbL | | GO:0030255 protein secretion by the type IV secretion system | GO:0005886 plasma membrane |
| | TRAF_ECOLI | Protein TraF | | | GO:0042597 periplasmic space |

### 3 GO term annotations of representative plasmid accessory proteins

| Function | Entry name | Protein name | GO terms (Molecular Function) | GO terms (Biological Process) | GO terms (Cellular Component) |
|---|---|---|---|---|---|
| Antibiotic resistance | AADB1_KLEPN | 2"-aminoglycoside nucleotidyltransferase | GO:0008871 aminoglycoside 2"-nucleotidyltransferase activity<br>GO:0046872 metal ion binding | GO:0046677 response to antibiotic | |
| | VANA_ENTFC | Vancomycin/teicoplanin A-type resistance protein VanA | GO:0005524 ATP binding<br>GO:0008716 D-alanine-D-alanine ligase activity<br>GO:0046872 metal ion binding | GO:0008360 regulation of cell shape<br>GO:0009252 peptidoglycan biosynthetic process<br>GO:0046677 response to antibiotic<br>GO:0071555 cell wall organization | GO:0005737 cytoplasm<br>GO:0005886 plasma membrane |
| Resistance to heavy metals | MERA_PSEAI | Mercuric reductase | GO:0016152 mercury (II) reductase activity<br>GO:0016668 oxidoreductase activity, acting on a sulfur group of donors, NAD(P) as acceptor<br>GO:0045340 mercury ion binding<br>GO:0050660 flavin adenine dinucleotide binding<br>GO:0050661 NADP binding | GO:0050787 detoxification of mercury ion | |
| | CADA1_STAAU | Cadmium-transporting ATPase | GO:0005524 ATP binding<br>GO:0008551 P-type cadmium transporter activity<br>GO:0016887 ATP hydrolysis activity<br>GO:0046872 metal ion binding | GO:0046686 response to cadmium ion | GO:0005886 plasma membrane |
| New metabolic process | HADB_CLODI | (R)-2-hydroxyisocaproyl-CoA dehydratase alpha subunit | GO:0016836 hydro-lyase activity<br>GO:0046872 metal ion binding<br>GO:0051539 4 iron, 4 sulfur cluster binding | GO:0006551 L-leucine metabolic process | |
| Virulence factors | CYAA_BACAN | Calmodulin-sensitive adenylate cyclase | GO:0004016 adenylate cyclase activity<br>GO:0005516 calmodulin binding<br>GO:0005524 ATP binding<br>GO:0008237 metallopeptidase activity<br>GO:0008294 calcium- and calmodulin-responsive adenylate cyclase activity<br>GO:0036094 small molecule binding<br>GO:0046872 metal ion binding<br>GO:0090729 toxin activity | GO:0006171 cAMP biosynthetic process<br>GO:0099004 calmodulin dependent kinase signaling pathway | GO:0005576 extracellular region<br>GO:0044164 host cell cytosol<br>GO:1902494 catalytic complex |

### 4 Comparison results of embeddings between ProtTrans and PlasGO

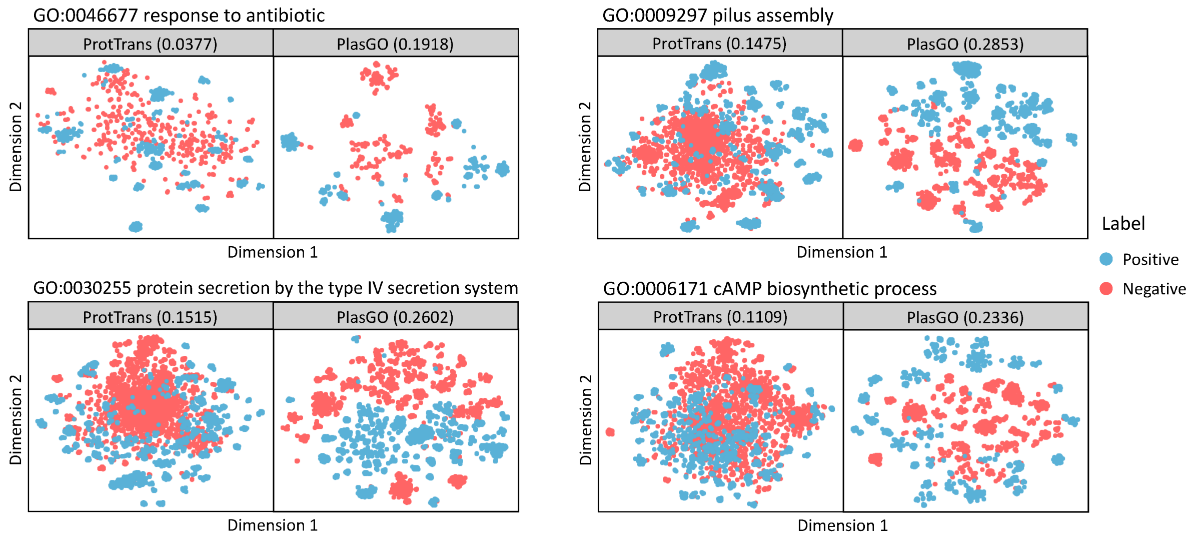


**Supplementary Figure S1.** Embedding comparisons for the BP binary classifications.

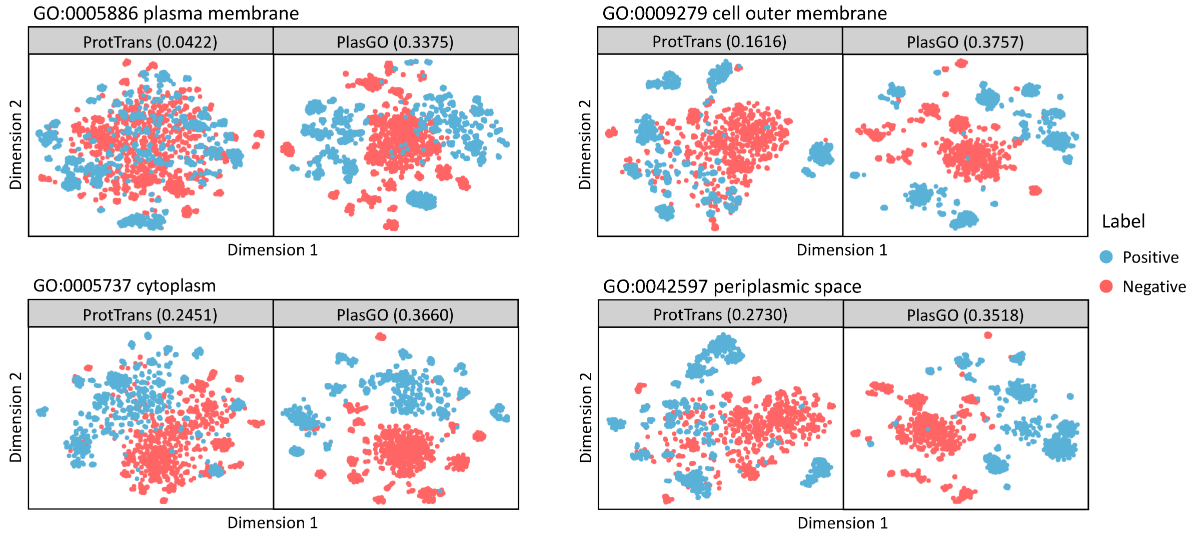


**Supplementary Figure S2.** Embedding comparisons for the CC binary classifications.

#### 5 Analysis of the elusive GO term labels

**Supplementary Table S1.** Detailed list of the elusive labels identified using the validation set. The third column represents the AUPR scores on the validation set for the elusive labels obtained from PlasGO, all of which are below 0.3. Furthermore, the fourth to seventh columns indicate the AUPR scores on the test set for the elusive labels obtained from PlasGO, PFresGO, TM-Vec, and DeepGOPlus (the top four tools), respectively. AUPR scores on the test set that exceed 0.3 are displayed in dark red.

| Category | GO term | PlasGO (val) | PlasGO | PFresGO | TM-Vec | DeepGOPlus | Detail |
|---|---|---|---|---|---|---|---|
| MF | GO:0004497 | 0.2303 | 0.8165 | 0.1933 | 0.7612 | 0.3892 | monooxygenase activity |
| MF | GO:0004659 | 0.0331 | 1.0 | 0.0857 | 0.3333 | 0.0001 | prenyltransferase activity |
| MF | GO:0016701 | 0.0338 | 0.0021 | 0.0018 | 0.0031 | 0.001 | oxygenase |
| MF | GO:0016765 | 0.0673 | 0.0818 | 0.0059 | 0.0222 | 0.0014 | transferase activity, transferring alkyl or aryl groups |
| MF | GO:0016805 | 0.0628 | 0.0321 | 0.0061 | 0.0239 | 0.0078 | dipeptidase activity |
| MF | GO:0016846 | 0.182 | 0.5 | 0.0004 | 0.0092 | 0.0001 | carbon-sulfur lyase activity |
| MF | GO:0019001 | 0.0563 | 0.0378 | 0.0541 | 0.041 | 0.0047 | guanyl nucleotide binding |
| MF | GO:0019114 | 0.0928 | 0.1535 | 0.1389 | 0.1285 | 0.0012 | catechol dioxygenase activity |
| MF | GO:0019205 | 0.1893 | 0.0757 | 0.1449 | 0.048 | 0.0006 | nucleobase-containing compound kinase activity |
| MF | GO:0030145 | 0.1062 | 0.0974 | 0.0608 | 0.0805 | 0.0228 | manganese ion binding |
| MF | GO:0030246 | 0.0227 | 0.2419 | 0.2145 | 0.0314 | 0.0055 | carbohydrate binding |
| MF | GO:0042910 | 0.1166 | 0.0004 | 0.0005 | 0.0004 | 0.0006 | xenobiotic transmembrane transporter activity |
| MF | GO:0043565 | 0.0474 | 0.0276 | 0.0109 | 0.0281 | 0.0168 | sequence-specific DNA binding |
| MF | GO:0046943 | 0.1303 | 0.0036 | 0.0015 | 0.0009 | 0.0003 | carboxylic acid transmembrane transporter activity |
| MF | GO:0051287 | 0.2041 | 0.1011 | 0.9431 | 0.2702 | 0.0232 | NAD binding |
| MF | GO:1901682 | 0.0333 | 0.0008 | 0.0011 | 0.0009 | 0.0007 | sulfur compound transmembrane transporter activity |
| BP | GO:0006081 | 0.0349 | 0.037 | 0.1425 | 0.1331 | 0.0352 | cellular aldehyde metabolic process |
| BP | GO:0006457 | 0.2902 | 0.1315 | 0.1132 | 0.0764 | 0.008 | protein folding |
| BP | GO:0007049 | 0.1268 | 0.0551 | 0.0672 | 0.0098 | 0.0088 | cell cycle |
| BP | GO:0009605 | 0.1261 | 0.0534 | 0.0396 | 0.043 | 0.0315 | response to external stimulus |
| BP | GO:0009607 | 0.0444 | 0.0435 | 0.0708 | 0.0383 | 0.0372 | response to biotic stimulus |
| BP | GO:0009628 | 0.0179 | 0.25 | 0.0011 | 0.0005 | 0.0001 | response to abiotic stimulus |
| BP | GO:0022402 | 0.0504 | 0.0196 | 0.0677 | 0.0084 | 0.0074 | cell cycle process |
| BP | GO:0042221 | 0.2663 | 0.0552 | 0.0363 | 0.0249 | 0.0184 | response to chemical |
| BP | GO:0042537 | 0.0963 | 0.3222 | 0.5608 | 0.0475 | 0.0093 | benzene-containing compound metabolic process |
| BP | GO:0042592 | 0.1638 | 0.0222 | 0.0054 | 0.0133 | 0.0042 | homeostatic process |
| BP | GO:0043603 | 0.2406 | 0.0332 | 0.0066 | 0.0065 | 0.0019 | amide metabolic process |
| BP | GO:0044419 | 0.0903 | 0.1196 | 0.1367 | 0.0623 | 0.0434 | interspecies interaction |
| BP | GO:0046451 | 0.0656 | 0.0452 | 0.2275 | 0.0181 | 0.0016 | diaminopimelate metabolic process |
| BP | GO:0048518 | 0.0178 | 0.0014 | 0.0026 | 0.0125 | 0.0008 | positive regulation of biological process |
| BP | GO:0048519 | 0.0815 | 0.0201 | 0.0225 | 0.0182 | 0.0155 | negative regulation of biological process |
| BP | GO:0048583 | 0.1985 | 0.0001 | 0.0002 | 0.0004 | 0.0001 | regulation of response to stimulus |
| BP | GO:0048878 | 0.1586 | 0.0245 | 0.007 | 0.0132 | 0.0042 | chemical homeostasis |
| BP | GO:0051172 | 0.0791 | 0.0189 | 0.0185 | 0.0184 | 0.0155 | inhibition of nitrogen metabolic process |
| BP | GO:0051301 | 0.0973 | 0.0761 | 0.0249 | 0.0063 | 0.0088 | cell division |
| BP | GO:0080134 | 0.2026 | 0.0002 | 0.0001 | 0.0004 | 0.0001 | regulation of response to stress |
| BP | GO:1901700 | 0.1417 | 0.0921 | 0.0408 | 0.0257 | 0.0184 | response to oxygen-containing compound |
| CC | GO:0009986 | 0.242 | 0.025 | 0.0955 | 0.0468 | 0.0285 | cell surface |
| CC | GO:0043226 | 0.0372 | 0.117 | 0.0629 | 0.0569 | 0.0622 | organelle |
| CC | GO:0043229 | 0.0661 | 0.1099 | 0.0747 | 0.0462 | 0.0615 | intracellular organelle |

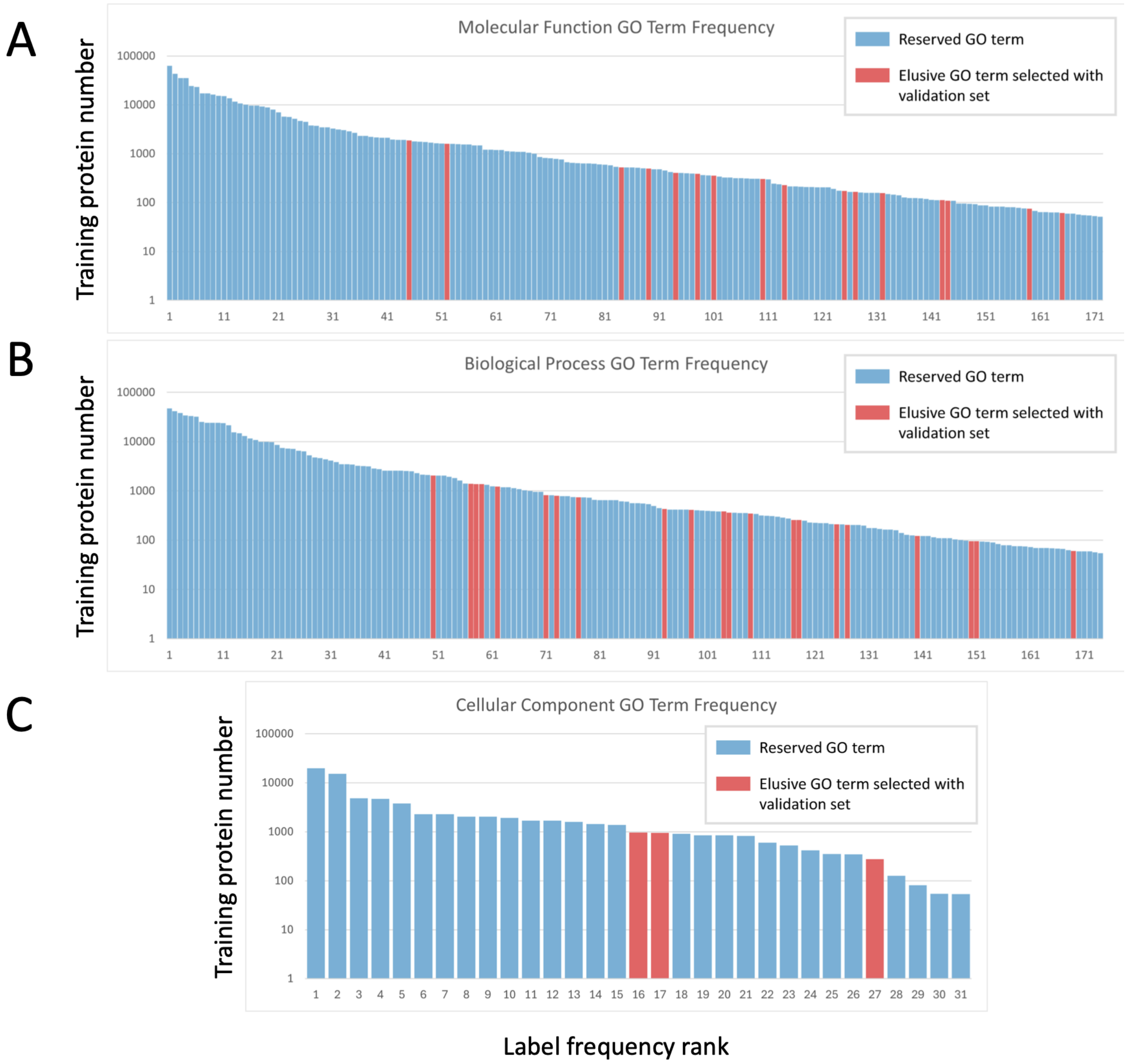


**Supplementary Figure S3.** Sorted occurrence frequency of GO term labels in the training set across three GO categories. The x-axis represents the ranks of the GO term labels based on their frequency, while the y-axis represents the frequency in exponential format (base 10). The red bars indicate the elusive labels, while the blue bars represent the remaining labels. It can be observed that most of the elusive labels are rare classes.

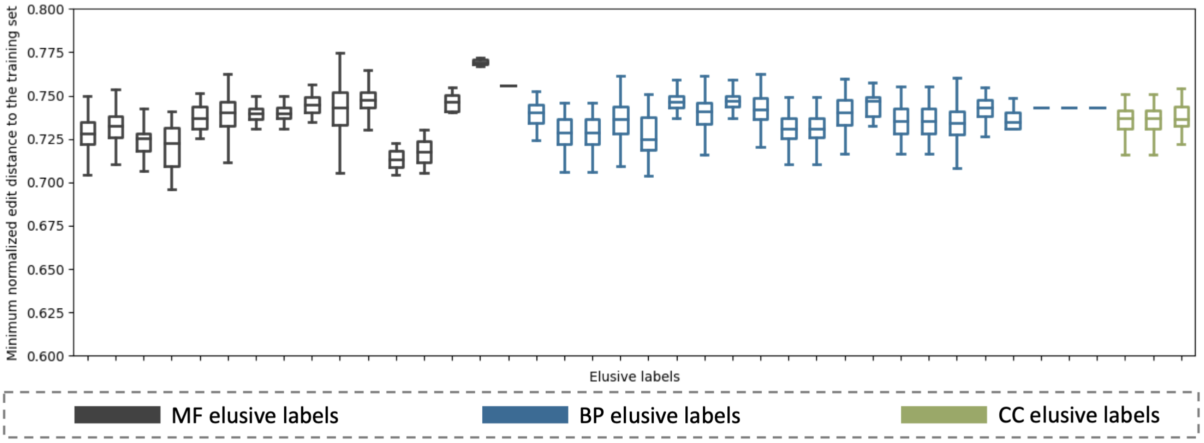


**Supplementary Figure S4.** The distribution of the distance between the training set and the test set for each elusive label. The distance distribution was measured by the minimum edit distance between each testing protein and the training set, normalized by dividing by the length of the longest sequence in the corresponding protein pair. The black, blue, and green boxes represent MF, BP, and CC labels, respectively. Furthermore, within each GO category, the elusive labels are sorted by their occurrence frequency in the training set. We can observe that all the minimum distances exceed 67.5%, indicating a low sequence similarity between the training set and test set for each elusive label [1].

**Supplementary Table S2.** The performance of different classification methods on the elusive labels evaluated using the RefSeq test set.

| Method | GO category | Fmax | AUPR |
|---|---|---|---|
| 3-layer DNN classifier | MF | 0.1237 | 0.1308 |
| | BP | 0.1578 | 0.0449 |
| | CC | 0.735 | 0.0314 |
| PlasGO (no context) | MF | 0.342 | 0.1787 |
| | BP | 0.1578 | 0.0439 |
| | CC | 0.735 | 0.0766 |
| PlasGO (standard) | MF | 0.3773 | 0.1983 |
| | BP | 0.1578 | 0.0677 |
| | CC | 0.735 | 0.084 |

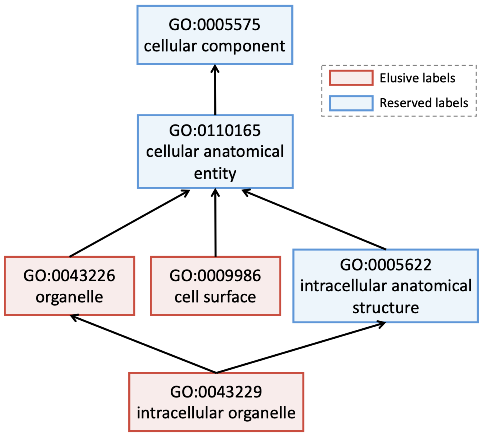


**Supplementary Figure S5.** The directed acyclic graph (DAG) structure including the three CC elusive labels (shown as red boxes) and their ancestor terms (shown as blue boxes).

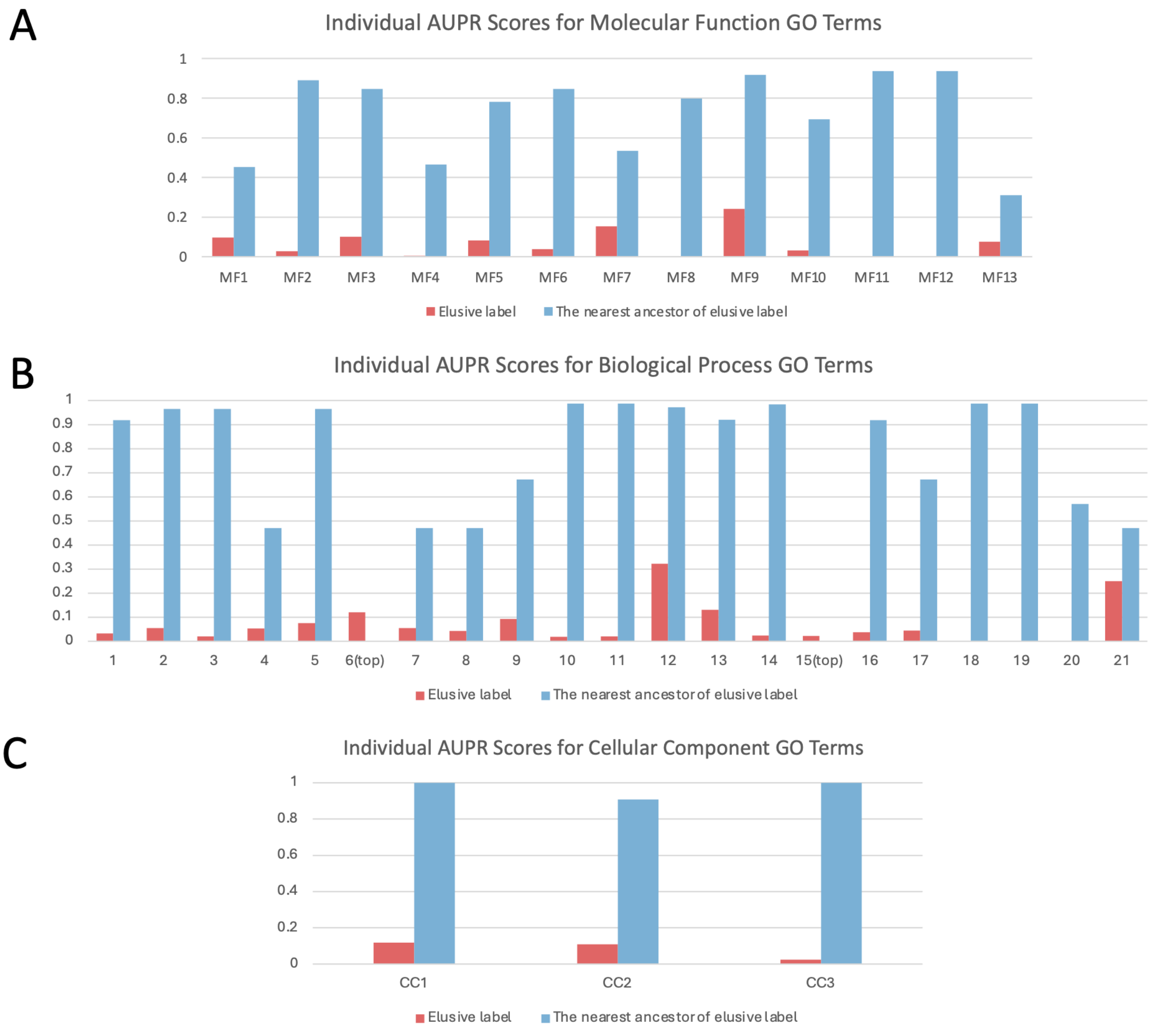


**Supplementary Figure S6.** The performance comparisons between the elusive labels and their nearest ancestor terms for the three GO categories. Notably, the three MF elusive labels that exhibited poor performance on the validation set but good performance on the test set, achieving AUPR scores of 0.8165, 1.0, and 0.5, respectively, are not shown. Besides, a suffix '(top)' is added to two out of the BP elusive labels, which indicates that they do not have reserved ancestor terms.

### 6 Evaluation of the learned confidence scores for PlasGO

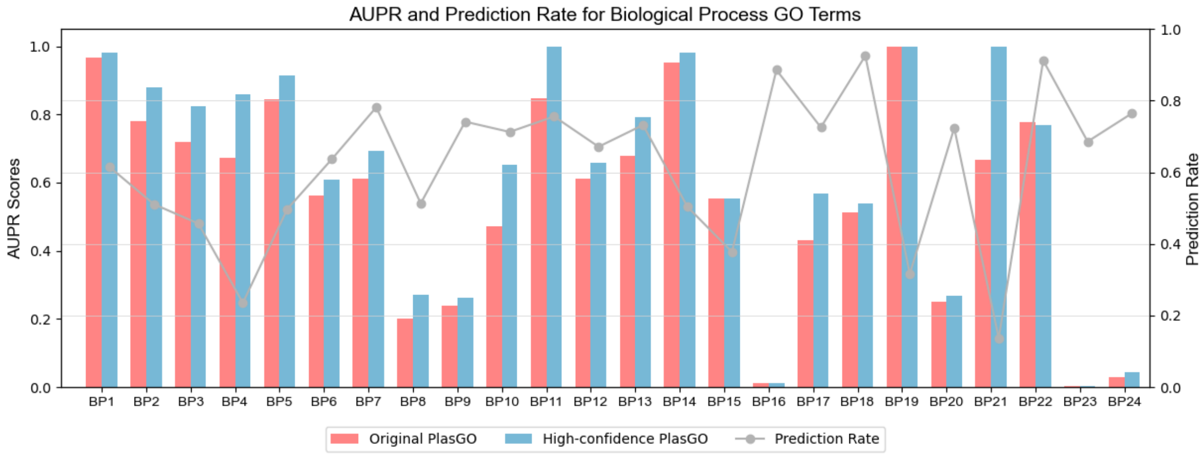


**Supplementary Figure S7.** The AUPR comparisons on the BP category between the original PlasGO and the high-confidence mode of PlasGO.

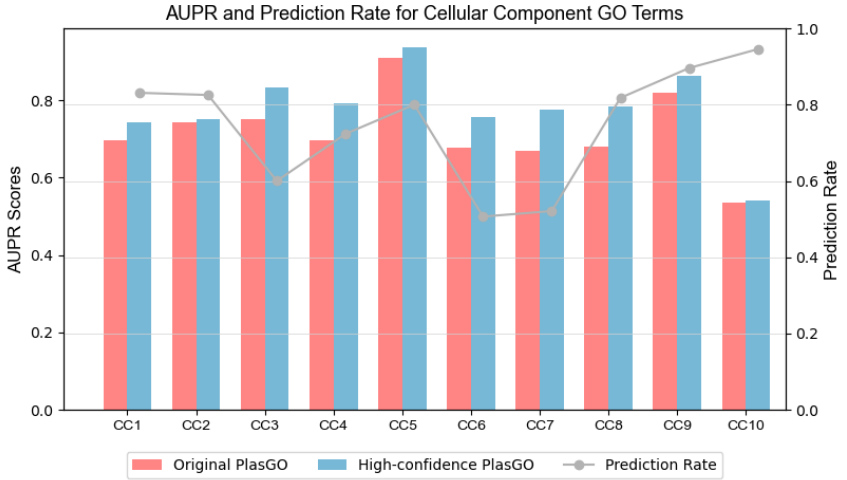


**Supplementary Figure S8.** The AUPR comparisons on the CC category between the original PlasGO and the high-confidence mode of PlasGO.

### 7 Distributions of the number of high-confidence predicted GO terms for unannotated proteins

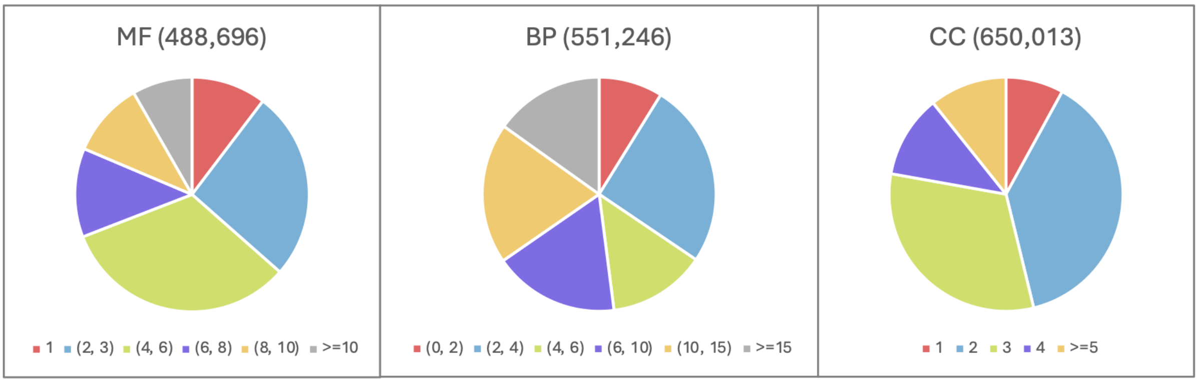


### 8 Complete list of the GO term indicators for the three core functions

| Core function | GO term | Detail |
|---|---|---|
| Replication | GO:0006260 | DNA replication |
| | GO:0003697 | single-stranded DNA binding |
| | GO:0003678 | DNA helicase activity |
| Stability | GO:0006276 | plasmid maintenance |
| | GO:0042802 | identical protein binding |
| | GO:0004519 | endonuclease activity |
| | GO:0004527 | exonuclease activity |
| Conjugation | GO:0009292 | horizontal gene transfer |
| | GO:0003917 | DNA topoisomerase type I (single strand cut, ATP-independent) activity |
| | GO:0009297 | pilus assembly |
| | GO:0030255 | protein secretion by the type IV secretion system |

### 9 Detailed information of the proteins encoded in the two well-studied plasmids

| Plasmid | Index | Protein ID | Gene product annotation | Protein class |
|---|---|---|---|---|
| pOLA52 | 1 | WP_001067858 | IS6-like element IS26 family transposase | MGE genes |
| | 2 | WP_000027057 | broad-spectrum class A beta-lactamase TEM-1 | Payload |
| | 3 | WP_000677445 | type 3 fimbria minor subunit MrkF | Conjugation |
| | 4 | WP_012291466 | type 3 fimbria adhesin subunit MrkD | Conjugation |
| | 5 | WP_000813718 | type 3 fimbria usher protein MrkC | Conjugation |
| | 6 | WP_000820818 | type 3 fimbria chaperone MrkB | Conjugation |
| | 7 | WP_002916128 | type 3 fimbria major subunit MrkA | Conjugation |
| | 8 | WP_228261368 | IS1-like element IS1A family transposase | MGE genes |
| | 9 | WP_001293129 | H-NS family nucleoid-associated regulatory protein | Payload |
| | 10 | WP_000850859 | hemolysin expression modulator Hha | Payload |
| | 11 | WP_012291470 | type IA DNA topoisomerase | Replication |
| | 12 | WP_000717624 | TrbM/KikA/MpfK family conjugal transfer protein | Conjugation |
| | 13 | WP_000722603 | cag pathogenicity island Cag12 family protein | Payload |
| | 14 | WP_012291471 | type IV secretory system conjugative DNA transfer family protein | Conjugation |
| | 15 | WP_012291472 | P-type DNA transfer ATPase VirB11 | Conjugation |
| | 16 | WP_012291473 | VirB10/TraB/TrbI family type IV secretion system protein | Conjugation |

| | | | | |
|---|---|---|---|---|
| | 17 | WP_000783379 | TrbG/VirB9 family P-type conjugative transfer protein | Conjugation |
| | 18 | WP_000394613 | type IV secretion system protein | Conjugation |
| | 19 | WP_000796673 | type IV secretion system protein | Conjugation |
| | 20 | WP_000748128 | EexN family lipoprotein | Conjugation |
| | 21 | WP_000744202 | type IV secretion system protein | Conjugation |
| | 22 | WP_012291475 | VirB3 family type IV secretion system protein | Conjugation |
| | 23 | WP_000916182 | TrbC/VirB2 family protein | Conjugation |
| | 24 | WP_001446885 | transcription termination/antitermination | Payload |
| | 25 | WP_000539530 | MobP1 family relaxase | Conjugation |
| | 26 | WP_000757693 | DNA distortion polypeptide 1 | Conjugation |
| | 27 | WP_000220560 | type II toxin-antitoxin system RelE/ParE family toxin | Stability |
| | 28 | WP_000121743 | plasmid stabilization protein | Stability |
| | 29 | WP_001050931 | RepB family plasmid replication initiator protein | Replication |
| | 30 | WP_001675596 | DNA distortion polypeptide 3 | Conjugation |
| | 31 | WP_012291478 | ParA family protein | Stability |
| | 32 | WP_000051066 | plasmid partition protein ParG | Stability |
| | 33 | WP_000864788 | ParA family protein | Stability |
| | 34 | WP_000203199 | recombinase family protein | Payload |
| | 35 | WP_000609146 | DinQ-like type I toxin DqlB | Payload |
| | 36 | WP_272056275 | DinQ-like type I toxin DqlB | Payload |
| | 37 | WP_001067858 | IS6-like element IS26 family transposase | MGE genes |
| | 38 | WP_063102497 | bleomycin binding protein | Payload |
| | 39 | WP_000084745 | pyridoxamine 5'-phosphate oxidase family protein | Payload |
| | 40 | WP_001067858 | IS6-like element IS26 family transposase | MGE genes |
| | 41 | WP_002914189 | multidrug efflux RND transporter periplasmic adaptor subunit OqxA | Payload |
| | 42 | WP_000888203 | Rrf2 family transcriptional regulator | Payload |
| pSK41 | 1 | WP_011117677 | YolD-like family protein | Payload |
| | 2 | WP_001273859 | recombinase family protein | Payload |
| | 3 | WP_000331763 | ArdC family protein | Stability |
| | 4 | WP_001252101 | MobA/MobL family protein | Conjugation |
| | 5 | WP_001819633 | parM protein | Stability |
| | 6 | WP_000358311 | recombinase | Payload |
| | 7 | WP_001008213 | helix-turn-helix transcriptional regulator | Payload |
| | 8 | WP_000043161 | replication initiator protein A | Replication |
| | 9 | WP_001106022 | IS6-like element IS257 family transposase | MGE genes |
| | 10 | WP_102695945 | protein rep | Replication |
| | 11 | WP_012695373 | sulfite exporter TauE/SafE family protein | Payload |
| | 12 | WP_001106022 | IS6-like element IS257 family transposase | MGE genes |
| | 13 | WP_227992126 | protein rep | Replication |
| | 14 | WP_000119405 | MobV family relaxase | Conjugation |
| | 15 | WP_001242578 | bleomycin binding protein | Payload |
| | 16 | WP_001795128 | aminoglycoside O-nucleotidyltransferase ANT(4')-Ia | Payload |
| | 17 | WP_001106019 | IS6-like element IS257 family transposase | MGE genes |
| | 18 | WP_000368849 | conjugative transfer protein TrsA | Conjugation |
| | 19 | WP_000591622 | CagC family type IV secretion system protein | Conjugation |
| | 20 | WP_000979860 | TrsD/TraD family conjugative transfer protein | Conjugation |
| | 21 | WP_000735569 | TrsH/TraH family protein | Conjugation |
| | 22 | WP_001094109 | DNA topoisomerase III | Replication |
| | 23 | WP_000209436 | type IV secretory system conjugative DNA transfer family protein | Conjugation |
| | 24 | WP_011117679 | conjugal transfer protein TrbL family protein | Conjugation |
| | 25 | WP_000608970 | single-stranded DNA-binding protein | Replication |
| | 26 | WP_011117680 | IS6-like element IS257 family transposase | MGE genes |
| | 27 | WP_001579520 | protein rep | Replication |
| | 28 | WP_001146389 | quaternary ammonium compound efflux SMR transporter QacC | Payload |
| | 29 | WP_001105984 | IS6-like element IS257 family transposase | MGE genes |
| | 30 | WP_000393259 | GNAT family N-acetyltransferase | Payload |
| | 31 | WP_001028144 | aminoglycoside O-phosphotransferase APH(2")-Ia | Payload |
| | 32 | WP_223200496 | IS6 family transposase | MGE genes |
| | 33 | WP_000889978 | type I toxin-antitoxin system Fst family toxin | Stability |
| | 34 | WP_001106006 | IS6-like element IS257 family transposase | MGE genes |
